# miRglmm: a generalized linear mixed model of isomiR-level counts improves estimation of miRNA-level differential expression and uncovers variable differential expression between isomiRs

**DOI:** 10.1101/2024.05.03.592274

**Authors:** Andrea M. Baran, Arun H. Patil, Ernesto Aparicio-Puerta, Marc K. Halushka, Matthew N. McCall

## Abstract

MicroRNA-seq data is produced by aligning small RNA sequencing reads of different miRNA transcript isoforms, called isomiRs, to known microRNAs. Aggregation to microRNA-level counts discards information and violates core assumptions of differential expression (DE) methods developed for mRNA-seq data. We establish miRglmm, a DE method for microRNA-seq data, that uses a generalized linear mixed model of isomiR-level counts, facilitating detection of miRNA with differential expression or differential isomiR usage. We demonstrate that miRglmm outperforms current DE methods in estimating DE for miRNA, whether or not there is significant isomiR variability, and simultaneously provides estimates of isomiR-level DE.

## Background

MicroRNA (miRNA) are a class of small, noncoding RNAs that perform a role in transcriptional regulation. They are typically 18-24 nucleotides long single-stranded RNA molecules that bind to mRNA causing translational suppression or mRNA degradation (1, 2). Through this mechanism, miRNA can regulate entire pathways and drive disease pathogenesis (3, 4). The specificity of each miRNA:mRNA interaction leads to discrete downstream consequences (5). Small RNA sequencing (sRNA-seq) can be used to measure miRNA expression. This process involves enriching samples for small RNA species prior to sequencing, followed by aligning the sequence reads to known miRNAs, tRNA fragments, or other RNA species (6). miRNA count data, which we refer to as miRNA-seq data throughout, is produced when sequence reads are aligned to known miRNAs, exclusively.

The specific mature sequence listed in miRNA databases is considered the canonical miRNA sequence (7). Different isoforms of individual miRNAs, called isomiRs, can arise from both variation in the nucleotide sequence or from variation in the transcript length (8). Sequence length variants have more or fewer nucleotides at the 5’ and/or the 3’ end of the canonical sequence, whereas polymorphic (internal) isomiRs include different nucleotides within the mature sequence (7). A third form of isomiR is the addition of an adenosine or uracil tail by a terminal uridylyl transferase (TUT) or similar enzyme (9). We will use the general term isomiR to describe the various sequence isoforms mapped to a miRNA, without distinction between biologically or technically derived isomiRs, with the assumption that in biological samples most sequences observed at sufficient levels to model are biologically relevant isomiRs.

The collection of isomiRs that align to a given miRNA are referred to as the miRgroup (10). As an example of isomiR diversity within miRgroup, Supplemental Tables 1 and 2 display isomiR-level miRNA-seq read counts for two miRgroups: hsa-let-7a-5p and hsa-miR-26a-5p. Typically, read counts of each isomiR in a miRgroup are aggregated by summation and summarized as a single read count for each miRNA (as shown in the last row of Supplemental Tables 1 and 2). There are numerous combinations of isomiR counts that produce the same miRNA counts; therefore, this aggregation leads to the loss of any information contained in individual isomiR expression.

IsomiRs are thought to possess unique biological roles (11). Like canonical sequences, isomiRs are conserved throughout evolution, and biogenesis of isomiRs is tightly regulated. The region of heterogeneity between isomiRs may have unique implications for miRNA-mediated translation regulation (8). 5’ isomiRs have modifications at the 5’ end of the miRNA resulting from differential processing of paralogous pre-miRNA and can result in regulation of distinct target mRNA (12). 3’ isomiRs are more common and result from post-transcriptional trimming or tailing sequence modifications (13). These modifications can alter miRNA function by impacting target recognition or extent of target repression. Additionally, these modifications can impact the stability of the molecule as adenylation protects miRNAs from degradation, while uridylation is associated with increased degradation (7).

Naturally occurring isomiRs have been shown to play distinct roles in a variety of biological processes including cytokine expression, virus proliferation, apoptosis, and tumor progression (11). Differential expression (DE) at the isomiR-level has identified isomiRs with cancer-specific expression (14). The biological importance of isomiRs highlights the need for a miRNA-seq analysis method that can account for distinct isomiR expression patterns in estimating miRNA-level differential expression, while also producing more granular estimates of isomiR-level differences.

Similar to differential expression analyses typically performed using bulk mRNA-seq data, the goal of most miRNA-seq studies is to study if, and which, miRNA differ in their expression between groups of samples. Common DE tools developed for mRNA-seq data include DESeq2 (15), edgeR (16) and limma-voom (17). All of these methods have been frequently used to analyze miRNA-seq data; however, there are some key differences between miRNA-seq data and mRNA-seq data that may make the assumptions of these methods invalid when applied to miRNA-seq data. Due to the compositional nature of RNA sequencing data, the reads can be viewed as a random sample of a fixed size from the pool of all RNA in the library, which can be modeled by a multinomial distribution. In bulk mRNA-seq data, the number of unique mRNA expressed is large and the reads are distributed relatively evenly across the mRNA. In this setting, the true underlying multinomial distribution can be approximated by marginal binomial distributions. This approximation is a key assumption of the tools mentioned above but is violated in miRNA-seq data in two important ways. First, there are generally fewer than 500 miRNA compared to over 10,000 mRNA expressed in a sample, which makes it more likely that random fluctuation in the expression of one miRNA substantially affects the expression of other miRNAs. Second, the distribution of reads in miRNA-seq data is often skewed toward a small number of highly expressed miRNA compared to the more uniform distribution of reads seen in mRNA-seq data (18). This can induce negative correlation between highly expressed miRNA (due to competition of being counted) regardless of their underlying biological correlation. This also results in sparse data with a small number of non-zero features making traditional normalization approaches, such as size factor normalization or counts-per-million (CPM) normalization, unstable or highly skewed.

Differences between miRNA and mRNA make the analysis of miRNA counts at the isomiR level feasible. First, even using Illumina short read sequencing, miRNAs and all isomiRs are fully sequenced (18-24 nt), whereas typically only fragments of mRNAs (100s of nt out of ∼5,000-50,000 nt species) are sequenced. Therefore, quantifying the expression of miRNAs based on the entire sequence is possible (6). Second, as previously stated, there are far fewer miRNA expressed compared to mRNAs. Maintaining and analyzing sequence level data at the scale needed for mRNA analysis would be computationally burdensome. Recent work has highlighted the importance of analyzing miRNA-seq data at the isomiR level (10). Through the analysis of 28 public miRNA-seq datasets and a newly generated human endothelial cell hypoxia data set, the authors show substantial differences between isomiR expression and their corresponding canonical miRNAs when applying DESeq2 to miRNA counts versus isomiR counts. As noted above, aggregate miRNA level data violates the assumption of independence between features due to the small number of unique miRNAs expressed in a sample and the highly skewed distribution of miRNA expression. While analyzing isomiR level data greatly increases the feature space and reduces the overall skew of the expression distribution, the isomiR level data introduce a new source of dependence due to high correlation between isomiRs from the same miRNA, which also violates a core assumption of the DESeq2 model. Despite this limitation, we agree with the authors’ conclusion that “a comprehensive re-evaluation of the miRNA-seq analysis practices” is needed.

Negative binomial mixed models (NBMM) can be used to model overdispersed count data when there is a correlation structure among the counts (19). This is the case when working with miRNA-seq data at the isomiR level, as counts that are observed for the same isomiR or for the same sample are correlated. Modelling the raw (non-normalized) counts directly has advantages over the typical counts per million (CPM) normalization. CPM is a dependent normalization strategy, so a change in any one miRNA read count will lead to changes in all other miRNA values even in the absence of a change in absolute expression (20). To account for the technical artifact of difference in the number of total reads between samples, one can incorporate an offset term into the NBMM to adjust for total overall read count within each sample. NBMMs have been proposed for other next-generation sequencing analyses, including RNA-seq, metagenomic sequencing, and single cell RNA-seq analyses (21–24). However, the NBMMs used in these methods all model aggregate feature level data rather than isoform level data and none have been applied to miRNA-seq data.

In this manuscript, we propose miRglmm, a method to model isomiR-level counts using a generalized linear mixed model to estimate miRNA-level DE while also obtaining estimates of isomiR DE and variability. We compare miRglmm to several commonly used DE tools on simulated data, an experimental benchmark data set, and real biological data sets. In both simulations and experimental benchmark data, miRglmm has a lower Mean Squared Error (MSE) than other DE tools and better confidence interval coverage. Additionally, we find significant isomiR-level variability exists within most miRgroups in real biological data sets, further motivating the use of miRglmm to analyze miRNA-seq data.

## Results

The methodology developed in this manuscript was motivated by two initial observations. First, different isomiRs of the same miRNA can behave very differently between groups of samples. To illustrate this, we selected a study of bladder and testes samples (25), with the goal of minimizing possible technical variation, and therefore capturing true biologically relevant isomiR-level differences. We observed isomiRs with uniformly zero counts within bladder and large non-zero counts in testes, and other isomiRs with uniformly zero counts within testes and large non-zero counts in bladder (Supplemental Table 1). Aggregation to miRNA counts masks these isomiR differences and results in a loss of information. Additionally, we observe evidence of differential isomiR usage between bladder and testes (Figure 1). We consider the canonical sequence (which is typically the most highly expressed sequence) and the next two highest expressing isomiRs, as these would contribute most to the aggregate miRNA count. Even in this small subset of isomiRs, we observe differences in DE between tissues across isomiRs, indicating the need for a model that can capture this variability in isomiR DE.

**Figure 1:**
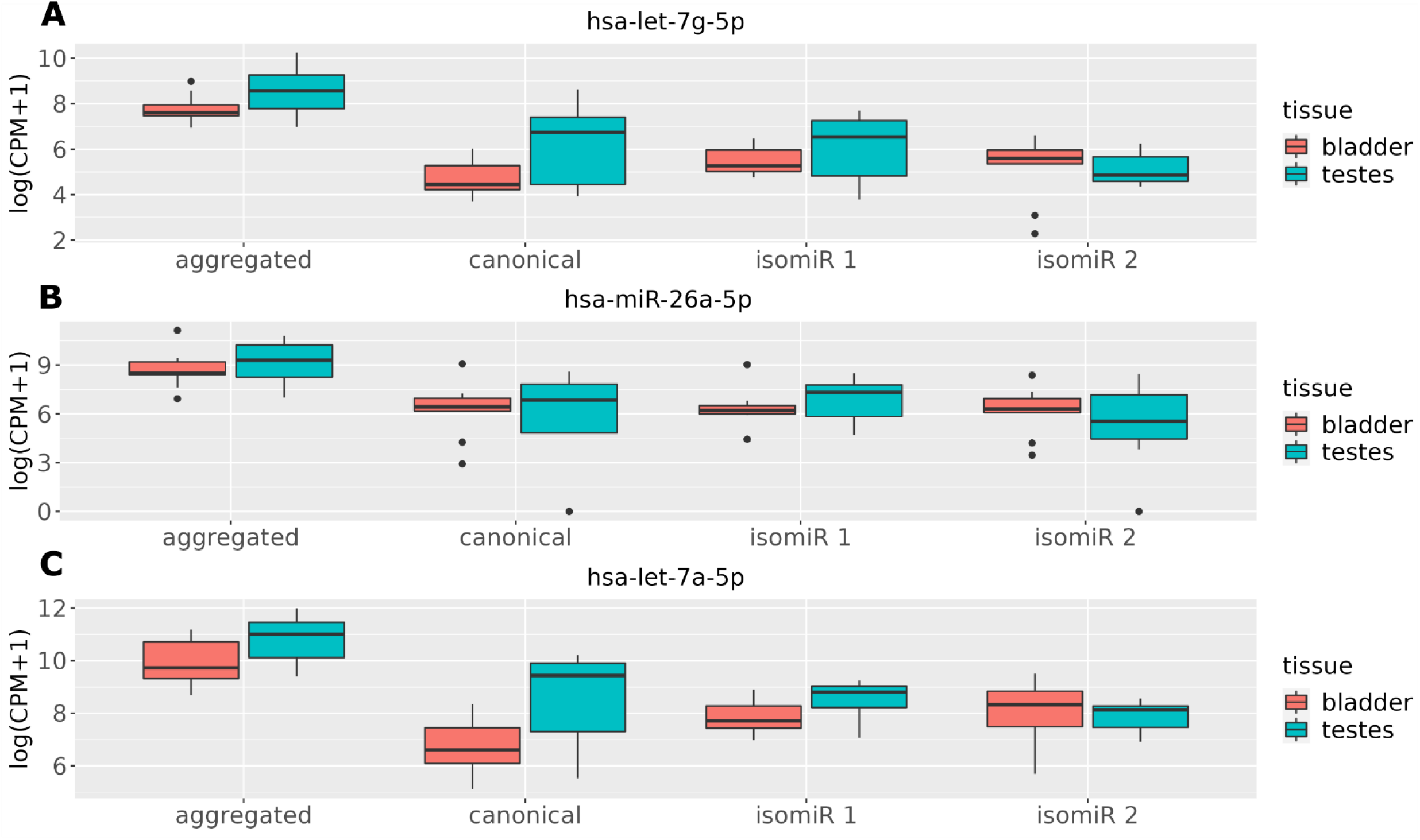
Tissue-specific counts are summarized by boxplots for three miRNAs (panel A: hsa-let-7g-5p, panel B: hsa-miR-26a-5p, panel C: hsa-let-7a-5p). Aggregated miRNA-level counts from summing counts across isomiRs within a miRNA are compared to the canonical/representative sequence and the two highest expressed non-canonical isomiRs (isomiR 1 and isomiR 2). The canonical sequence counts, and isomiR 1 are all higher in testes than bladder, but isomiR 2 has the opposite trend, which is masked when counts are aggregated to miRNA-level. CPM: counts per million.

Second, while miRNA-level data exhibit an artificially induced negative correlation between highly expressed miRNAs, isomiR-level data do not always exhibit this bias. Data generated by high-throughput RNA sequencing is fundamentally compositional in nature. In other words, the resulting reads can be viewed as a random sample of a fixed size from the pool of RNA generated during library preparation. This produces a competition-to-be-counted (26) in which randomly measuring more of one feature decreases the chance of measuring other features. Consider the extreme example of only two features; with a fixed number of total reads, it is clear that a higher count for feature 1 implies a lower count for feature 2, resulting in perfect negative correlation between the two features regardless of their underlying biological correlation. Frequently, this induces a negative correlation between the two most highly expressed miRNAs; however, this induced correlation is not always observed for the most highly expressed isomiRs. To illustrate this, we selected a study of immune cell types (27), with the goal of analyzing samples without cellular heterogeneity that can occur in tissue-level samples. We compared the correlation between the two highest expressed miRNAs with the correlation between the two highest expressed isomiRs for each cell type. While we observed a negative correlation between the highest expressed miRNAs for all cell types, the same was not true at the isomiR level where we observed a mix of positive and negative correlations depending on the cell type (Supplemental Figure 1). This indicates that utilizing the isomiR-level count data could overcome technical biases seen with aggregated miRNA-level count data.

### miRglmm: a generalized linear mixed effects model for miRNA-seq data analysis

We developed a method and corresponding software to analyze isomiR-level counts from miRNA-seq data. Our method, miRglmm, accounts for dependencies introduced by reads coming from the same sample and from the same isomiR, by using a negative binomial generalized linear mixed model with random effects for sequence and sample. We utilized a negative binomial model to directly model the counts without the need for transformation and include an offset term to normalize for sequencing depth (see Methods for details). Unlike existing methods, miRglmm provides estimates of differential expression at both the miRNA-level and the isomiR-level. miRglmm is implemented in a free and open-source R package available at: https://github.com/mccall-group/miRglmm.

### miRglmm outperforms other methods in detecting differentially expressed miRNAs in the presence of isomiR variability

To assess the performance of miRglmm in comparison to existing methods (DESeq2 (15), edgeR (16), limma-voom (17) and Negative Binomial Generalized Linear Models (NB GLM)(28)), we used a collection of monocyte samples to simulate differential expression at both the miRNA-and isomiR-level (see Methods). Over 100 simulations, miRglmm provided the lowest mean MSE (mean = 0.015) and median MSE (median = 0.014) among all the methods (Table 1, Supplemental Table 3). miRglmm was also the method with the lowest standard deviation across simulations (standard deviation (SD) = 0.005) and the lowest maximum MSE (maximum= 0.033). miRglmm minimized MSE in 97% of simulations (97/100) when comparing methods within a given simulation. DESeq2 minimized the MSE in the remaining 3 simulations. On average, for these 3 simulations, the MSE for DESeq2 was only smaller than the MSE for miRglmm by 0.001. miRglmm provided the highest mean coverage proportion (mean = 0.91) and highest median coverage proportion (median = 0.93) among the 4 methods that produce standard error (SE) estimates which can be used to calculate confidence interval coverage (Table 1, Supplemental Table 4).

**Table 1:** MSE and 95% confidence interval coverage proportion summary statistics across 100 simulations.

| method | mean<br>MSE ($10^{-3}$) | minimum<br>MSE ($10^{-3}$) | maximum<br>MSE ($10^{-3}$) | # of times<br>method<br>minimizes<br>MSE | mean<br>coverage<br>proportion | minimum<br>coverage<br>proportion | maximum<br>coverage<br>proportion |
| --- | --- | --- | --- | --- | --- | --- | --- |
| miRglmm | 14.85 | 7.84 | 32.92 | 97 | 0.91 | 0.71 | 0.98 |
| DESeq2 | 21.92 | 12.29 | 36.08 | 3 | 0.80 | 0.66 | 0.89 |
| limmavoom | 23.10 | 13.81 | 45.37 | 0 | 0.79 | 0.55 | 0.89 |
| NB GLM | 23.15 | 13.26 | 48.83 | 0 | 0.77 | 0.53 | 0.89 |
| edgeR | 23.15 | 13.25 | 48.83 | 0 | NA | NA | NA |

MSE: Mean Squared Error, NA: not applicable

We can also assess performance within each “truth” group of miRNA: induced positive effect (N=20), induced negative effect (N=20), or no effect induced (N=82). When we calculate MSE within each group and summarize across simulations, we see that miRglmm provides markedly lower MSE in the groups with the induced effect compared to all aggregation methods (Figure 2A). In the group of miRNA with no change induced, DESeq2 tends to provide a lower MSE in most simulations. Similarly, the coverage proportion in the two groups of miRNA with induced effects tends to be much closer to the 95% nominal level than for the aggregation methods (Figure 2B). The aggregation methods have very poor confidence interval coverage proportions (0.3-0.4). For the miRNA with no induced effect, DESeq2 tends to have the highest coverage proportion, though miRglmm still provides coverage near the 95% nominal level.

**Figure 2:**
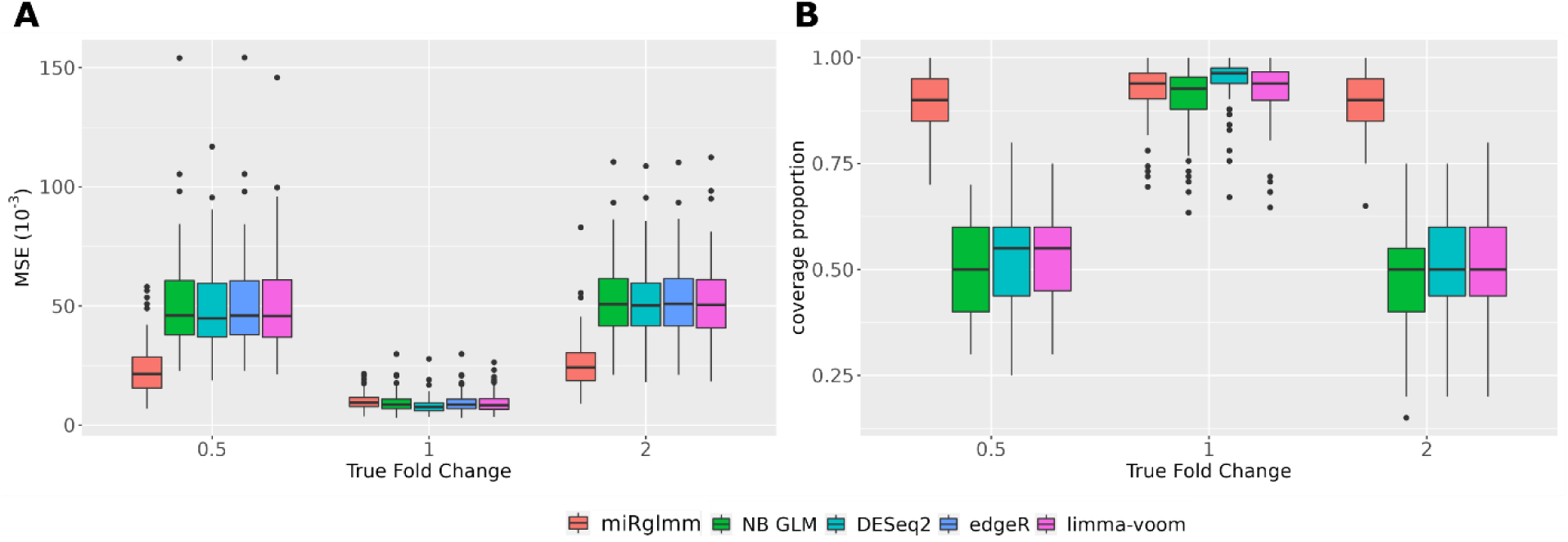
miRglmm outperforms the aggregation methods in terms of Mean Squared Error (MSE) for miRNA in which an effect is induced (panel A). DESeq2 provides the lowest MSE when there is no effect induced. The 95% confidence interval coverage proportion of miRglmm is much higher than the coverage proportion of the aggregation methods when an effect is induced (panel B). Results are based on 100 independent simulated data sets.

An additional advantage of miRglmm is that it provides an estimate of the variability in the group effect between isomiRs, facilitating the detection of miRNA with differential isomiR usage between groups. The other methods cannot estimate or account for this variability, and thus do not have the ability to detect miRNAs with differential isomiR usage. We observe that the proportion of miRNAs with significant variability between isomiRs is very high in the groups with the induced effect (Supplemental Figure 2), indicating that our simulation procedure correctly implemented random isomiR DE variability in the group effect. Taken together, these results support the conclusion that aggregation of isomiR counts to miRNA-level counts and the resulting loss of information is detrimental to performance in cases where significant isomiR DE variability exists. miRglmm, which can account for this variability, provides consistently high performance whether or not there is significant isomiR DE variability.

### miRglmm provides accurate estimates of differential expression for isomiRs

Along with providing miRNA-level estimates of differential expression, miRglmm also provides isomiR-level estimates of differential expression. While rarely done, DESeq2 can also provide isomiR-level estimates if run on the isomiR-level count matrix. Summarizing over 100 simulations, miRglmm (MSE=0.022) provides better estimates of isomiR-level differential expression than DESeq2 (MSE=0.041). When we assess performance within each “true effect” group, we see that miRglmm outperforms DESeq2 when there is a negative effect (MSE=0.035 vs 0.044), when there is a positive effect (MSE=0.035 vs 0.044), and when there is no differential expression (MSE=0.015 vs 0.040). This contrasts with what we observed at the miRNA-level, where miRglmm slightly underperformed DESeq2 with respect to MSE when there was no differential expression, but performed much better than DESeq2 when there was either positive or negative differential expression.

### miRglmm outperforms other methods in detecting differentially expressed miRNAs even when there is no isomiR variability

While variable isomiR usage is common in real biological data, we wanted to assess how miRglmm would perform in the absence of isomiR variability. To assess the performance of miRglmm in this context, we used a multi-protocol, multi-institution experimental benchmark data originally designed to compare the performance of four different sRNA-seq library preparation methods (29). This experiment used ratiometric pools of synthesized small RNAs with known variable amounts of differential expression (see Methods), providing a known true fold change value for each miRNA that can be used to evaluate the accuracy of DE methods. Because the data came from mixtures of synthesized miRNA pools, there should be no biological variability in isomiR usage.

We used MSE and confidence interval coverage to assess the performance of the DE methods (Table 2) as well as expression-based isomiR filtering thresholds (Supplemental Table 5). miRglmm accurately estimated the known DE magnitudes between synthetic miRNA pools while maintaining greater than nominal confidence interval coverage. In terms of MSE, miRglmm at any isomiR expression filtering threshold provided smaller MSE than all aggregation methods. However, miRglmm with no filtering of lowly expressed isomiRs had the highest MSE, and we observed that the estimates were systemically biased towards the null due to the inclusion of very lowly expressed isomiRs (Supplemental Figure 3). The four aggregation methods provide similar estimates but underestimate the true effect at all levels.

**Table 2:** Comparing performance of miRglmm to aggregation methods.

| method | MSE ($\times 10^{-3}$) | coverage<br>proportion |
| --- | --- | --- |
| miRglmm | 7.81 | 0.98 |
| DESeq2 | 10.19 | 0.99 |
| NB GLM | 12.11 | 0.97 |
| edgeR | 12.11 | NA |
| limma-voom | 13.78 | 0.99 |

MSE: Mean Squared Error, NA: not applicable

We chose log(median CPM) > -1 as the default filter for miRglmm because this retained the most isomiRs while achieving similar MSE as more stringent filtering thresholds (Supplemental Figure 4). As filtering gets more restrictive, we lose the ability to model some miRNA if fewer than two isomiRs are retained.

To determine whether other methods would improve if we filtered lowly expressed isomiRs, we compared MSEs based on estimates from running the other methods on aggregated counts from all isomiRs versus filtering prior to aggregation of the isomiR counts. The gain in performance for miRglmm due to filtering was not seen for the aggregation methods (Supplemental Figure 3). Since aggregating is done via summation, the low counts do not influence the sum to the same degree that they influence the miRglmm estimate.

Due to the synthetic nature of the data, we do not expect any biological isomiRs and presume that the sequence isoforms we are modeling arise solely from technical variation. Hence, the random slope parameter was not found to be significant in the synthetic data (Supplemental Figure 3). If we remove this extraneous random slope parameter for isomiR from the model, the performance is similar in terms of the MSE, confidence interval coverage and bias (Supplemental Table 6 and Supplemental Figure 5). This indicates that retaining the random slope parameter, even if not needed, does not diminish the ability of miRglmm to estimate the effect of interest.

### Differential miRNA expression and differential isomiR usage between immune cell types

We performed differential expression analyses comparing miRNA expression between five immune cell types: monocytes (N=39), Natural Killer Cells (N=38), CD4+ T lymphocytes (N=35), CD8+ T lymphocytes (N=32) and CD19+ B lymphocytes (N=26). The samples were obtained from one experiment in the miRNAome data (27), limiting the possibility of technical variation due to library preparation or sequencer differences. Based on isomiR filtering strategies assessed using data from the Extracellular RNA Communication Consortium (ERCC), we chose to implement miRglmm using a threshold of log(median CPM) < -1 to remove isomiRs with low expression. After isomiR filtering, we retained 7,389 isomiRs from 104 miRNA, and the number of isomiRs mapping to any one miRNA ranged from 6 to 633 with a median of 35 isomiRs.

We calculated log fold-change (logFC) estimates for each miRNA using miRglmm and existing DE methods (see Methods). B lymphocytes were used as the reference group in each logFC estimate. We used intra-class correlation coefficients (ICC) to assess agreement in logFC estimates between methods (Figure 3A). While all methods provided similar logFC estimates (ICC ≥ 0.95), we observed less agreement regarding statistical significance (Bejamini Hochberg False Discovery Rate (FDR) of 0.05) between methods (Cohen’s Kappa 0.36-0.53). There does not appear to be substantial differences in ICC between contrasts, i.e. specific cell type comparisons (Supplemental Figures 6-9), but some contrasts have very poor agreement in terms of statistical significance (minimum Kappa=0.11 when comparing edgeR and miRglmm for CD4+ T lymphocyte vs B lymphocyte and CD8+ T lymphocyte vs B lymphocyte contrasts). An upset plot of statistically significantly DE miRNAs across all contrasts provides a visual comparison of all 5 methods simultaneously (Figure 3B). There are 4 miRNA-contrast pairs significant for miRglmm only but no aggregation method (Supplemental Table 7), and 26 miRNA-contrast pairs significant for all aggregation methods but not miRglmm (Supplemental Table 8). We used likelihood ratios tests (LRTs) to assess if there is variability in the cell type effect between isomiRs within a given miRNA. All miRNAs had significant LRT p-values for the random slope parameter for isomiRs (Figure 3C). The other methods we assessed cannot estimate or account for this variability.

**Figure 3:**
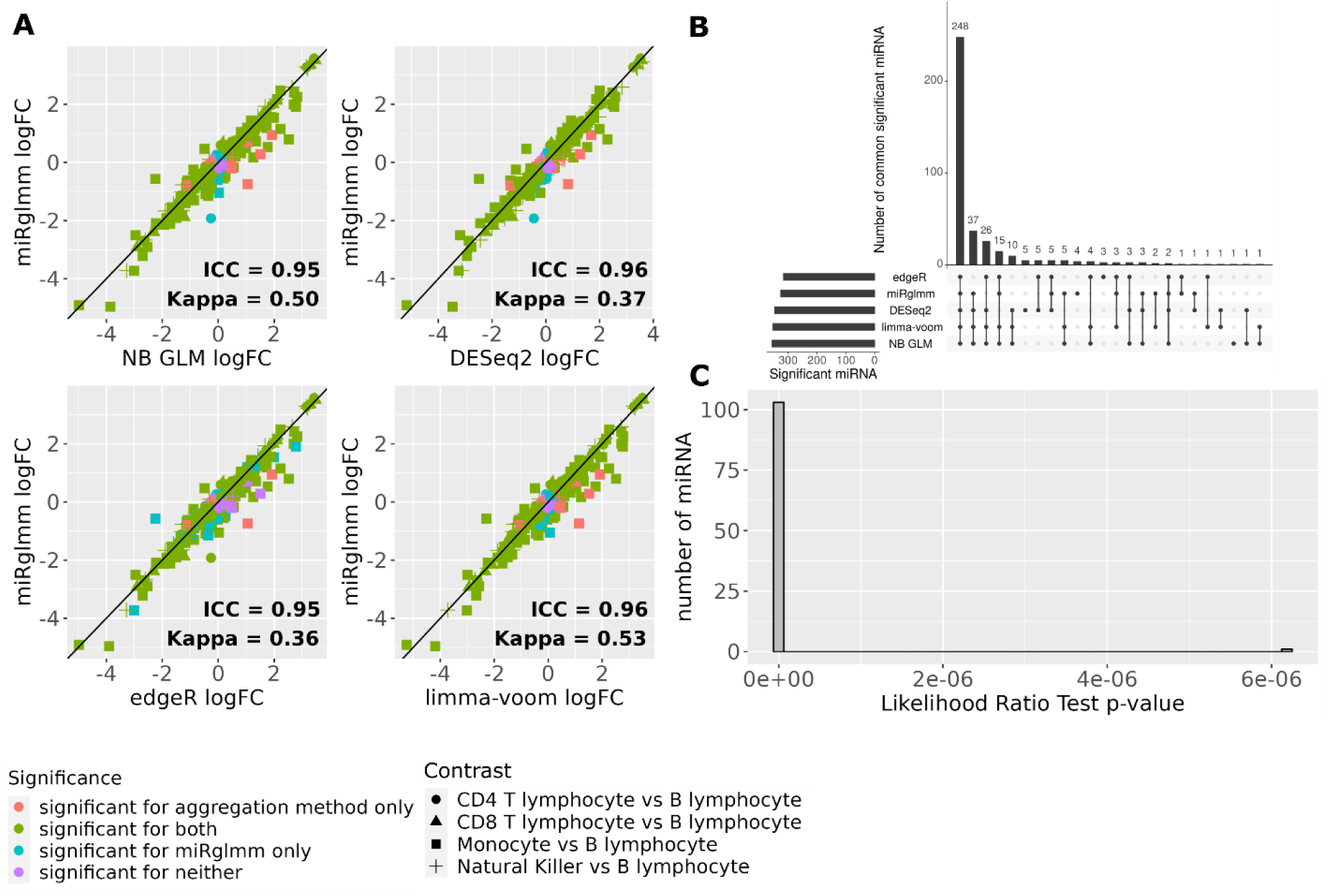
Log fold change (logFC) estimates of miRNA-level expression in cell types compared to B lymphocytes from miRglmm are plotted versus logFC estimates from each aggregation method (panel A). Each point is a unique contrast for a unique miRNA. Significant DE vs B lymphocytes is indicated by the color (Benjamini Hochberg FDR of 0.05). Intra-class correlation coefficients (ICC) are used to assess agreement in the values of the estimated logFC between methods. Cohen’s kappa is used to assess agreement in the significance of the estimate between methods. An upset plot shows the common number of miRNAs differentially expressed across the 5 methods (panel B). A histogram of the p-values of an LRT testing the significance of the random isomiR cell-type effect indicates that most of the miRNA have significant variability in the cell-type effect across isomiRs (panel C).

### Differential miRNA expression and differential isomiR usage between tissues

We performed differential expression analyses comparing miRNA expression between bladder (N=9) and testes (N=7). These tissue types represented the largest set of two distinct tissues in one single experiment in the miRNAome data. Since the data came from one experiment, there was no known technical source of variation to adjust for, and tissue-type was the only fixed effect in the model. We performed miRNA-level and isomiR-level filtering (see Methods), and retained 3,036 isomiRs from 90 miRNA. The number of isomiRs mapping to any one miRNA ranged from 5 to 436 with a median of 18 isomiRs.

We estimated log fold-change (logFC) between tissue types for each miRNA using miRglmm and compared the estimates to existing DE methods (see Methods). We used intra-class correlation coefficients to assess agreement in logFC estimates between methods (Figure 4A). All methods provided reasonably similar logFC estimates (ICC ≥ 0.8). As with the cell type analysis, agreement with respect to statistically significant DE, measured via Cohen’s Kappa, was much weaker. We similarly compared all aggregation methods in a pairwise fashion (Supplemental Figure 10) and observed high ICC (≥ 0.88) and weak-to-moderate kappas (≥ 0.35-0.68) between methods. An upset plot of miRNA deemed significantly different between bladder and testes across the methods provides a visual comparison of all 5 methods simultaneously (Figure 4B). 67 of 90 miRNA have significant LRT p-values for the random tissue parameter for isomiRs (Figure 4C).

**Figure 4:**
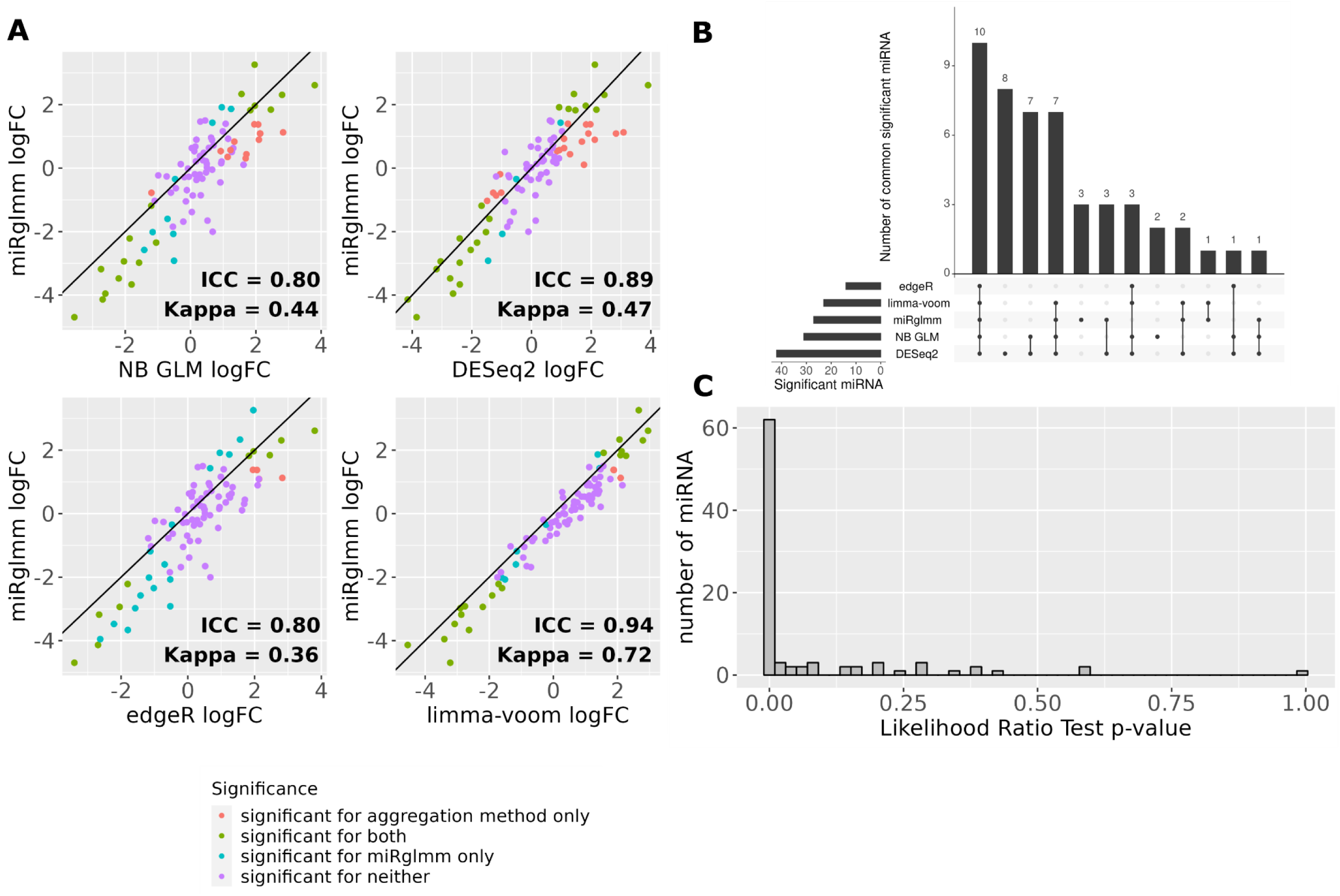
Log fold change (logFC) estimates of miRNA-level expression in testes compared to bladder from miRglmm are plotted versus logFC estimates from each aggregation method(panel A). Each point is a unique miRNA. Significant DE is indicated by the color (Benjamini Hochberg FDR of 0.05). Intra-class correlation coefficients (ICC) are used to assess agreement in the values of the estimated logFC between methods. Cohen’s kappa is used to assess agreement in the significance of the estimate between methods. An upset plot shows the common number of miRNAs significantly different between bladder and testes across the 5 methods (panel B). There are 3 miRNA found to be significant for all aggregation methods but not miRglmm. There are 3 miRNA found to be significant for miRglmm but none of the aggregation methods. A histogram of the p-values of an LRT testing the significance of the random isomiR tissue effect indicates that most of the miRNA have significant variability in the bladder-testes effect across isomiRs (panel C).

miRglmm can also provide isomiR-level estimates of differential expression. We visualized isomiR-level expression and variability within several miRNA by plotting the estimated effects for each isomiR mapping to the miRNA (Figure 5). We overlaid estimated fixed effects from miRglmm to compare the miRNA-level differential expression estimates (fixed effect estimates from miRglmm) with underlying individual isomiR effects (random isomiR effect estimates from miRglmm). We also overlaid the NB GLM estimated effect as an example aggregation method (other aggregation method results are similarly shown in Supplemental Figure 11). Figure 5A-5C show the 3 miRNA that only miRgImm identifies as differentially expressed between bladder and testes (Figure 4B). Figure 5D-5F show the 3 miRNA that are found to be significantly differentially expressed by all aggregation methods but not by miRglmm (Figure 4B). miRglmm estimates of expression tend to follow patterns observed for the majority of isomiRs, whereas NB GLM estimates of expression tend to follow patterns observed in only the highest expressing isomiRs. In the case of hsa-miR-143-5p (Figure 5B) there is little variability in isomiRs, increasing the precision of the estimate which results in a statistically significant difference when using miRglmm but not for NB GLM where the estimated effect is similar but less precise. All miRNA represented in Figure 5 with the exception of hsa-miR-143-5p (Figure 5B) were found to have significant random slope variability.

**Figure 5:**
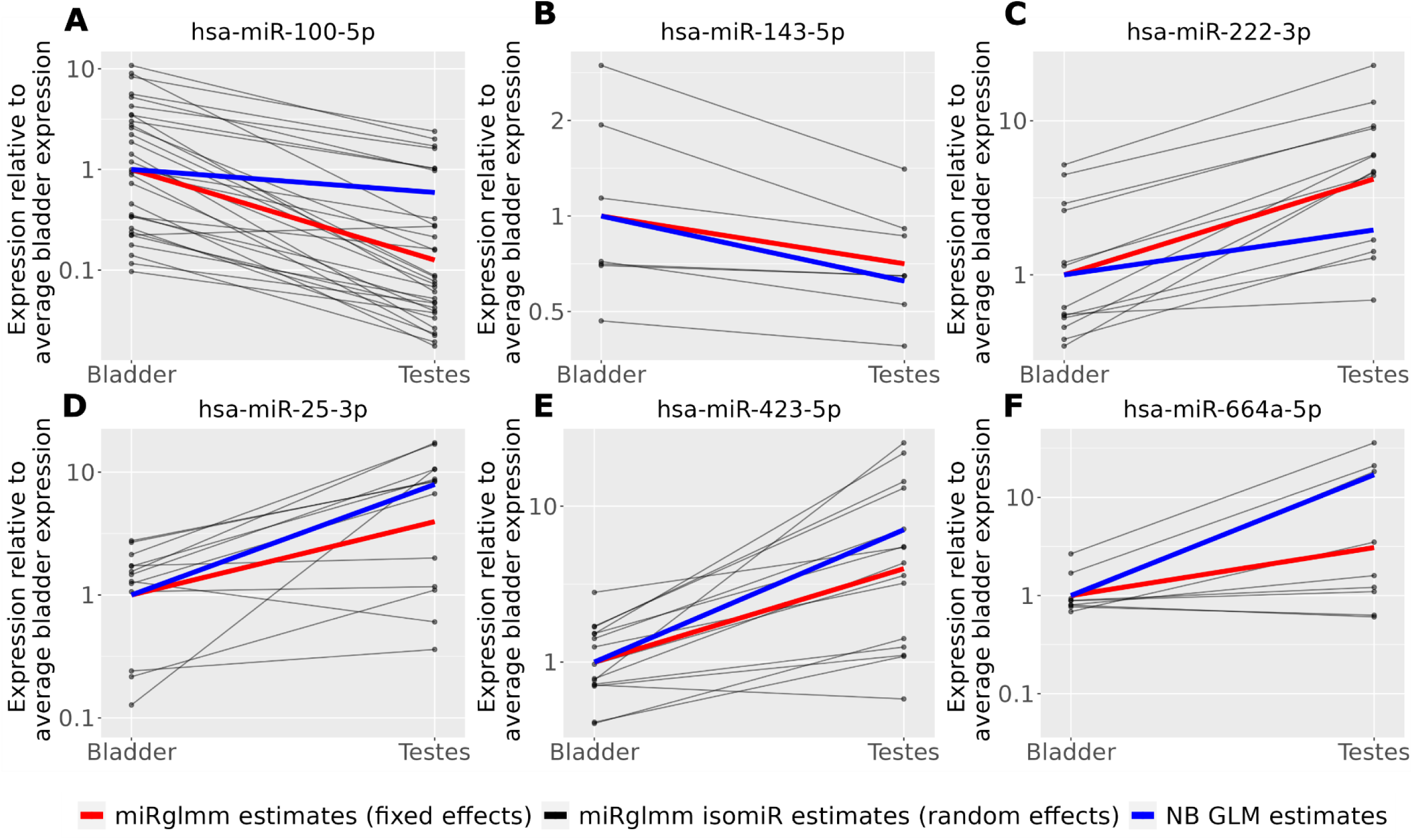
miRglmm estimates of relative expression are compared to estimates of relative expression from aggregated-count NB GLM models. We obtain estimates of isomiR variability within miRNA from the random effects components of miRglmm. Panels A-C represent miRNA found to be differentially expressed using miRglmm but not found by any of the aggregation methods. Panel D-F represent miRNA found to be differentially expressed using all aggregation methods but not found by miRglmm. With the exception of hsa-miR-143-5p (panel B), all miRNA presented have significant isomiR variability in the bladder vs testes effect.

## Discussion

With miRglmm, we address the need for a DE tool specifically suited for miRNA-seq data that can also utilize information from isomiRs in estimating miRNA-level DE estimates (10). Commonly used DE tools, such as DESeq2 (15), edgeR (16) and limma-voom (17), were built for analysis of mRNA-seq data, and some key features of miRNA-seq violate the independence between features assumption of these methods. Additionally, these tools run on aggregated miRNA-level counts, resulting in the loss of information contained in individual isomiRs. There is evidence of the biological importance of isomiRs (11), so miRglmm allows for the discovery of heterogeneous DE among isomiRs of the same miRNA via estimation of the random group effect for isomiR (and associated likelihood ratio test).

We show that miRglmm can identify miRNA with significant isomiR variability. When such variability exists, miRglmm far outperforms other DE tools in terms of MSE and confidence interval coverage, indicating that miRglmm provides a better estimate of DE at the miRNA-level than those produced by other DE tools. Even in settings where isomiR variability does not exist, performance of miRglmm remains superior to other DE tools, albeit to a lesser degree. Consistent isomiR expression reduces variability and leads to a precise estimate of miRNA-level differential expression. miRglmm also produces isomiR-level estimates that can be used for exploration of individual isomiR effects.

A limitation of miRglmm is that it does not aim to distinguish a true biological isomiR from a technically arising isomiR and treats all isomiRs within a miRgroup equally. It has been shown that differences in library preparation can induce bias in quantification of isomiRs (30, 31). By filtering lowly expressed isomiRs, we eliminate some technical variability due to background count alignment. Our study designs in the DE analysis of immune cell types as well as bladder vs testes tissue were implemented to minimize technical variation by selecting samples processed by a single laboratory using consistent library preparation techniques and sequencing equipment. Importantly, we observed the majority of miRNA did have significant isomiR variability between cell types / tissues in both data sets where we assume we are capturing primarily biological variability. If we were to analyze larger datasets that encompass multiple studies, we would expect even higher rates of miRNA with significant isomiR variability, though some of this would be purely technical. In this case, miRglmm is flexible and can adjust for technical design variables, such as laboratory, library preparation method, or sequencer, as we did in analyzing the ERCC benchmark data set (29).

## Conclusions

We provide a new method and analysis tool, miRglmm, that uses isomiR variability to improve differential expression analysis of miRNAs from miRNA-seq datasets. miRglmm provides superior performance to alternative DE tools, whether or not significant isomiR variability exists, and estimates both miRNA-level and isomiR-level differential expression.

## Methods

### miRglmm: A Negative Binomial GLMM-based DE tool for miRNA-seq data

The statistical model implemented in miRglmm is as follows. Let *C_ij_* denote the count for sample *i* = 1,…,*n* and isomiR *j =* 1,…,*J*. Let *T_i_* = ∑*^J^_j_*_=1_ *C_ij_* be the total counts for sample *i*. This is used as an offset term in the model to adjust for variable sequencing depth across samples. For each sample *i*, let *X_i_* be a *p*-vector of covariates. For each miRNA, miRglmm fits the following negative binomial mixed model (NBMM):

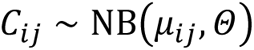

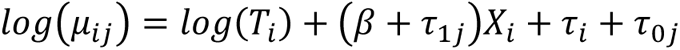

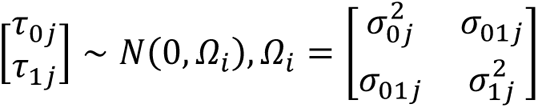

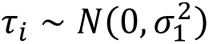

*β* is the fixed effect of primary interest. *τ*_*i*_ is the random intercept term for sample *i*, *τ*_0*j*_ is the random intercept for isomiR *j*, and *τ*_1*j*_ is the random slope term for isomiR *j*. The random effects for sample and isomiR are independent, while the intercept and slope random effects for isomiR have a dependence structure specified by *Ω*. The variance of the random slope *σ*^2^ can be used to assess variable DE between isomiRs. We built miRglmm using the function glmer.nb in the lme4 package, which fits an NBMM via a Laplace approximation to the maximum likelihood with a variety of optimizer choices (32). The analyses presented in this paper are reproducible using code in the GitHub repository found at https://github.com/mccall-group/miRglmm-paper.

### Software, data structures, and outputs

miRglmm is an R library that consists of one core function that can be easily integrated into DE analysis pipelines by replacing DE methods designed for mRNA-seq (such as DESeq2, edgeR, limma/voom).

IsomiR-level count matrices in the form of core Bioconductor structures, SummarizedExperiment objects, are taken as input, and a list of model fit summaries for each miRgroup analyzed is returned. The function can be run in parallel across miRgroups and can handle flexible design matrices. Additional functions are included in the package to extract and summarize values of interest from the model fit summaries, including miRNA-level estimates of DE with confidence intervals, isomiR-level estimates of expression within a miRgroup, estimates of variability in DE across isomiRs within a miRgroup, and a likelihood ratio test for significant isomiR variability within a miRgroup. The miRglmm package vignette includes examples of how to import isomiR-level count data from either miRge (33) or sRNAbench (34).

### The microRNAome data resource

The miRNAome dataset was assembled to more fully understand miRNA expression patterns across primary cell types (35, 36). The dataset was built upon 2,077 samples from 175 public datasets across 196 primary cell types. miRNA annotation and quantification was performed using the miRge3.0 pipeline (33). Briefly, miRge3.0 is a multi-step miRNA alignment program. From a FASTQ file, miRge3.0 collapses identical sequences and processes them through repeated Bowtie alignment steps to identify canonical miRNAs, isomiRs, and other RNA species. In addition to a final read count per miRNA, it outputs an alignment file containing counts of all aligned isomiRs, which was used here.

### Filtering of lowly expressed miRNAs and isomiRs

Counts per million (CPM) normalization of aggregated counts was used to assess overall miRNA expression, with the goal of retaining miRNA with sufficient expression to model. A threshold of log(median CPM)>5 was used to retain miRNA for modelling. Even after filtering at the miRNA-level, the resulting count matrix contains very sparse isomiR-level counts so miRglmm also filters lowly expressed isomiRs that contribute low/no amount of information to the model. Specifically, CPM normalization of isomiR-level counts was used to assess isomiR expression, and an isomiR filter based on log(median CPM) is implemented as an input argument of miRglmm (default = -1).

### Inducing known effects in real biological data to simulate differential expression

We created ground truth data by inducing a known artificial effect into real biological data, allowing us to assess the performance of miRglmm under conditions seen in real data. We searched the miRNAome data (35) for one cell type with large sample counts coming from the same study to have a relatively homogenous starting dataset. We used 39 monocyte samples from one study (27) and retained 122 miRNA with sufficient expression to model using a log(median CPM) cutoff of 5, as described above.

To induce an artificial “group” effect, we randomly split the samples into 2 groups, with 19 samples labelled as Group A and 20 samples labelled as Group B (Supplemental Figure 12). Of 122 total miRNA, we used stratified sampling to select 20 miRNA to be overexpressed in Group A, and another 20 miRNA to be underexpressed in Group A. The sampling was stratified by total miRNA expression to manipulate miRNA across the full range of expression values. We used the default log(median CPM) cutoff of -1 to retain isomiRs for analysis. For the 20 miRNA overexpressed in Group A, we multiplied the counts of all retained isomiRs in the miRgroup by a random truncated normal variable with mean value of 2, variance of 1, lower bound of 1 and upper bound of 3 in Group A samples only. These 20 miRNAs now have a known miRNA-level fold change (B vs A) = 0.5, with random variability between isomiRs. For the 20 miRNA underexpressed in Group A, we multiplied the counts of all retained isomiRs in the miRgroup by a random truncated normal variable with the same parameters as above but this time in Group B samples only. These 20 miRNAs now have a known miRNA-level fold change (B vs A) = 2, with random variability between isomiRs. Importantly, we recalculated the total counts used in the offset term after the artificial signal is induced. The stratification used in sampling miRNA ensures the effect of the signal being added is consistent across groups, even though the total counts increase for all samples. The entire procedure, including randomly splitting samples into two groups, was repeated 100 times.

For each miRNA, we estimated a miRNA-level differential group effect, measured via log fold-change (logFC) estimates, using miRglmm. We compared these estimates to differential group effects produced by other commonly used differential expression tools used on aggregated data: DESeq2 (15), edgeR (16), and limma-voom (17). We also included a Negative Binomial Generalized Linear Models (NB GLM), fit using glm.nb from the MASS R package (28), which is similar to the miRglmm model but without random effects that can model the isomiR variability. Data was aggregated to the miRNA-level prior to running NB GLM, DESeq2, edgeR and limma-voom, and hence we collectively call these methods “aggregation methods”. Aggregated count values were produced by summing counts from all isomiRs that align to a given miRNA.

We used Mean Squared Error (MSE) and 95% confidence interval coverage proportions to assess the performance. The MSE compared the estimated logFC to the induced effect for each miRNA, and the coverage proportion assessed the proportion that the true logFC fell in the 95% confidence interval estimated by the model. We cannot estimate a confidence interval coverage proportion for edgeR as that algorithm does not provide standard error (SE) estimates. We also separately looked at each method’s MSE and coverage when the effect was induced up or down, and those where no effect was induced.

To identify miRNA with significant variability in the group effect between isomiRs, we tested whether the random slope parameter τ_1j_ in the miRglmm model contributes significant information with a 1-degree of freedom (1-df) Likelihood Ratio Test (LRT) comparing likelihoods from the model specified above and a model removing only the τ_1j_ parameter. With this test, we can identify miRNA with significant random slope effects and summarize by induced effect groups.

### Analysis of a benchmark experiment using synthetic miRNA pools

We used experimental benchmark data with known expression differences to assess the performance of miRglmm and existing methods to estimate DE. The Extracellular RNA Communication Consortium (ERCC) was established to facilitate expansion of the field of extracellular RNA (exRNA) biology and consisted of collaborative projects to develop robust methods for isolation and analysis of exRNA data (37). One of these projects was a multi-protocol, multi-institution assessment of the bias of four sRNA-seq library preparation methods, using ratiometric pools of synthesized small RNAs (29). Chemically synthesized RNA oligonucleotides were added in varying ratios to pool A and pool B, from 10 to 1 and 1 to 10, leading to 15 levels of DE (from logFC = -2.3 to logFC = 2.3), providing a known fold change value for each miRNA that can be used to evaluate the accuracy of miRglmm and other methods (Supplemental Figure 13).

The ERCC sequence runs with 4N method (4 random nucleotides on both ends of the reads) were chosen (n=104 runs). The reference sequence database consisting of 286 human miRNAs and 48 other spikein miRNAs were indexed using bowtie-index. The reads were processed for illumina adapters ’TGGAATTCTCGGGTGCCAAGGA’ using cutadapt (38) followed by alignment using bowtie aligner (39). The parameters for bowtie include "No mismatch (-n 0)", "Trim 4 nucleotides on both ends (-5 4 -3 4)", "Avoid alignment against the reverse-complement reference strand (--norc)", and output in SAM format (-S). The SAM files were processed for read counts across each mapped miRNA and to account for PCR duplicates (4N-based) using custom Python scripts. The miRNA counts were used for the downstream analysis in miRglmm.

The ERCC data was processed by 6 different laboratories (Lab), and we used dimensionality reduction using non-metric multidimensional scaling (NMDS) to assess if there was sufficient laboratory variability to require adjustment for Lab in the analysis. The samples separated along the first dimension by Lab, and the second dimension appeared to capture the Pool effect (Supplemental Figure 14A). We included a fixed effect for Lab in the models for all methods to adjust for the technical effect of the differences in lab-specific sample handling, processing, and sequencer on the counts. The ERCC dataset included synthetic RNAs that were much longer at 50-90 nucleotides than the typical 18-24 nucleotide miRNAs. We excluded any RNA specified to have a length > 45, resulting in a set of 303 small RNAs.

This synthetic dataset also provided justification for establishing filters at both the miRNA and isomiR level. Since the oligonucleotides were synthetically added, we expect all included miRNA to be expressed. The distribution of miRNA expression via log(median CPM) of aggregated counts in this data can be used to determine a suitable threshold for miRNA expression, supporting our choice of log(median CPM)>5 used in all analyses. To filter sequences, we aimed to separate biologically relevant isomiR counts from random background expression. Since the ERCC contains sequence isoforms that do not map to known miRNA, we can consider these background counts and compare the distribution of expression levels to isomiR counts that map to known miRNA to identify an appropriate range for an isomiR-level filter. When we compared the distribution of background sequence expression (reads not mapping to any known miRNA) to expression of sequences that map to miRNA, we saw that the distributions separated around a log(median CPM) value of -1 (Supplemental Figure 14B). We ran miRglmm without any sequence filtering, and also compared the performance of miRglmm under a range of reasonable filters (log(median CPM)>-1 (default) to 2). We also assessed the effect of filtering prior to aggregation when using the aggregation methods.

For each miRNA, we estimated a differential Pool effect, adjusted for Lab and measured via logFC estimates, using miRglmm under a variety of isomiR filters. We also aggregated the data to the miRNA count level and ran the aforementioned aggregation methods for comparison. We used MSE and 95% confidence interval coverage proportions to assess the performance as described above. We utilized LRT as previously described to assess if there was significant variability in the Pool effect between isomiRs.

### Differential expression analyses of real biological data of tissues and cell types

Our goal in selecting samples for these analyses was to minimize possible technical variation, and therefore capture true biologically relevant isomiR-level differences. We chose bladder and testes tissues to compare because they represented a large set of tissues available from a single study (25), where sample processing and sequencer would be consistent across samples. Additionally, we chose a set of immune cell types that were also derived from a single study as our cell type differential expression analysis set (27). We aimed to produce miRNA-level differential expression (DE) estimates using miRglmm and compare these results to other commonly used differential expression tools that run on aggregated miRNA-level count data.

For each miRNA, we estimated a miRNA-level differential group (tissue or cell type) effect, measured via log fold-change (logFC) estimates, using miRglmm. We compared these estimates to differential group effects produced by existing aggregation methods using an intraclass correlation coefficient (ICC).

Significance at a Benjamini Hochberg False Discovery Rate (FDR) of 0.05 was compared using Cohen’s Kappa. We also used an upset plot to simultaneously compare miRNA considered differentially expressed between all 5 methods. We can produce isomiR-level estimates of expression within group by summing fixed and random effects for each isomiR. We utilized LRT as previously described to identify miRNA within which there is significant variability in the group effect between isomiRs.

## Declarations

### Ethics approval and consent to participate

Not applicable

### Consent for publication

Not applicable

### Availability of data and materials

Biological sample metadata used in this manuscript were accessed through the microRNAome database (35, 36). The original raw count data for the specific studies analyzed can be accessed via the sequence read archive (SRA) under the following accession numbers: SRP110505 (27) and SRP007946 (25). Synthetic data used in this manuscript were accessed via SRA accession number SRP098949 (29). Processed data files used in this manuscript can be found on Zenodo (10.5281/zenodo.12571079).

### Authors’ contributions

MNM and MKH conceived the project. AMB and MNM analyzed data. MKH interpreted results. AMB, AP, EAP and MNM developed the software. AMB and MNM wrote the manuscript. All authors read, edited and approved the final manuscript.

## Acknowledgements

We acknowledge the helpful feedback and discussion from other members of the McCall research group.

---

**Supplemental Table 1:**
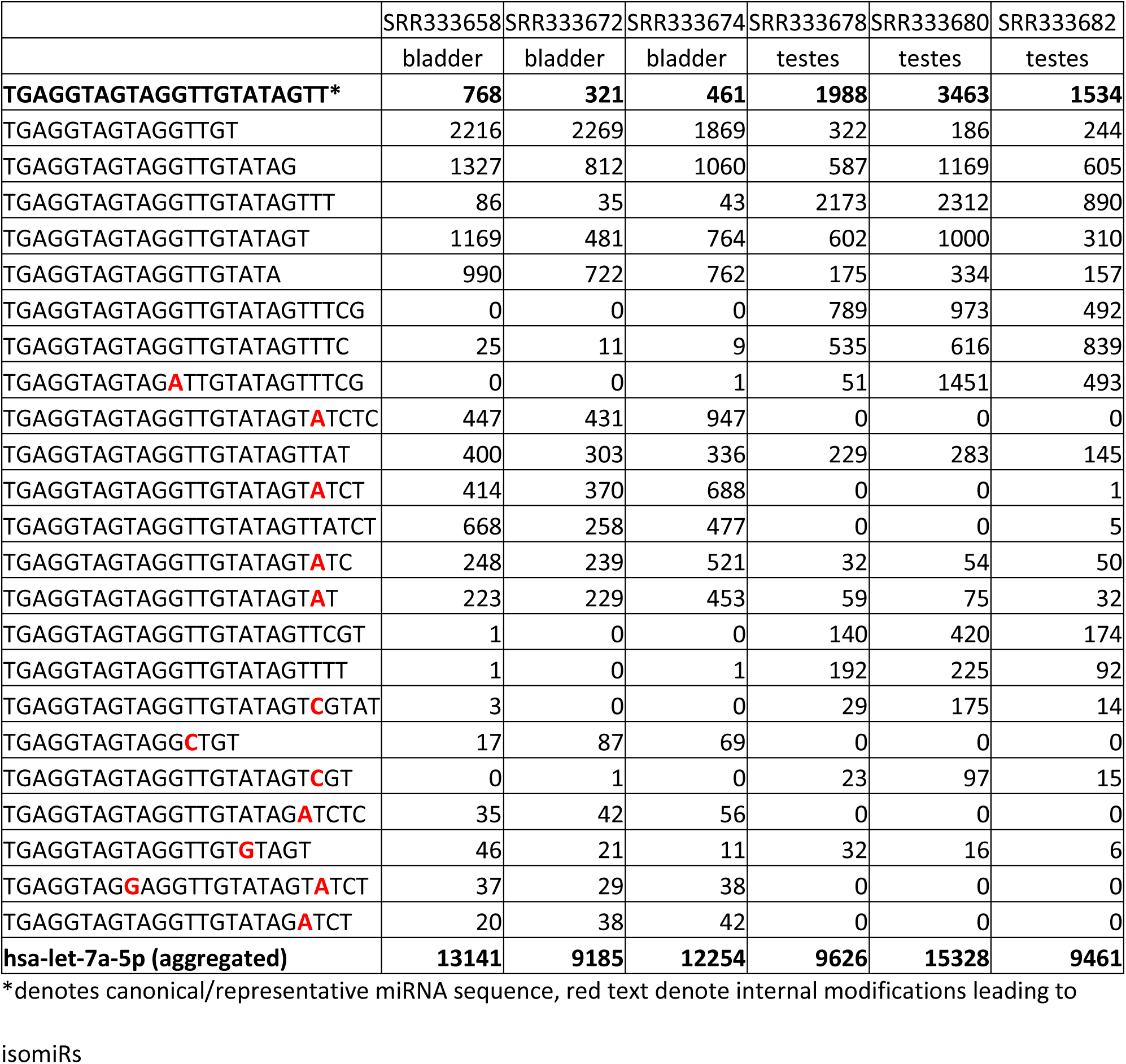
Example of isomiR data for miRNA hsa-let-7a-5p.

**Supplemental Table 2:**
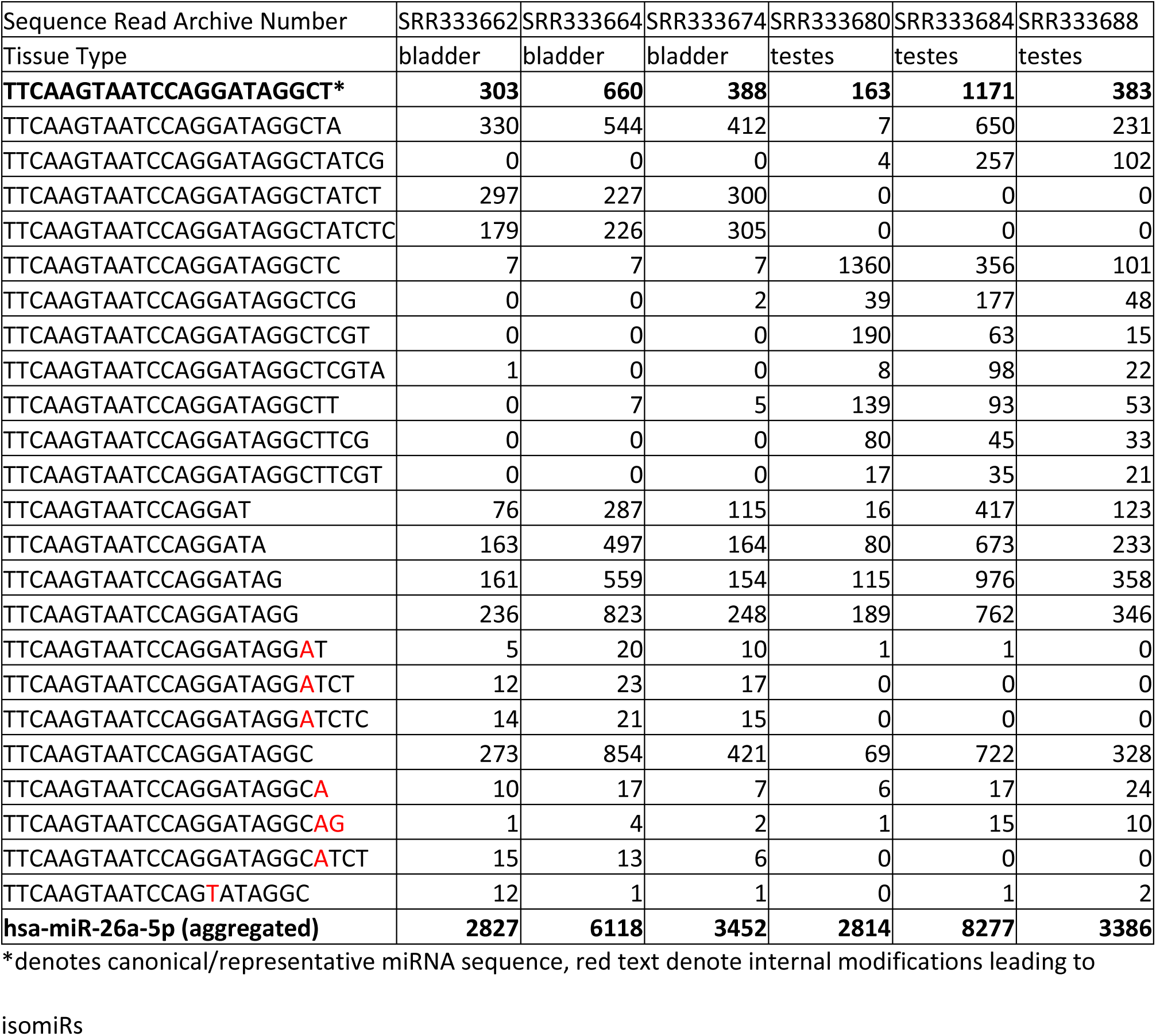
Example of isomiR data for miRNA hsa-miR-26a-5p.

**Supplemental Figure 1:**
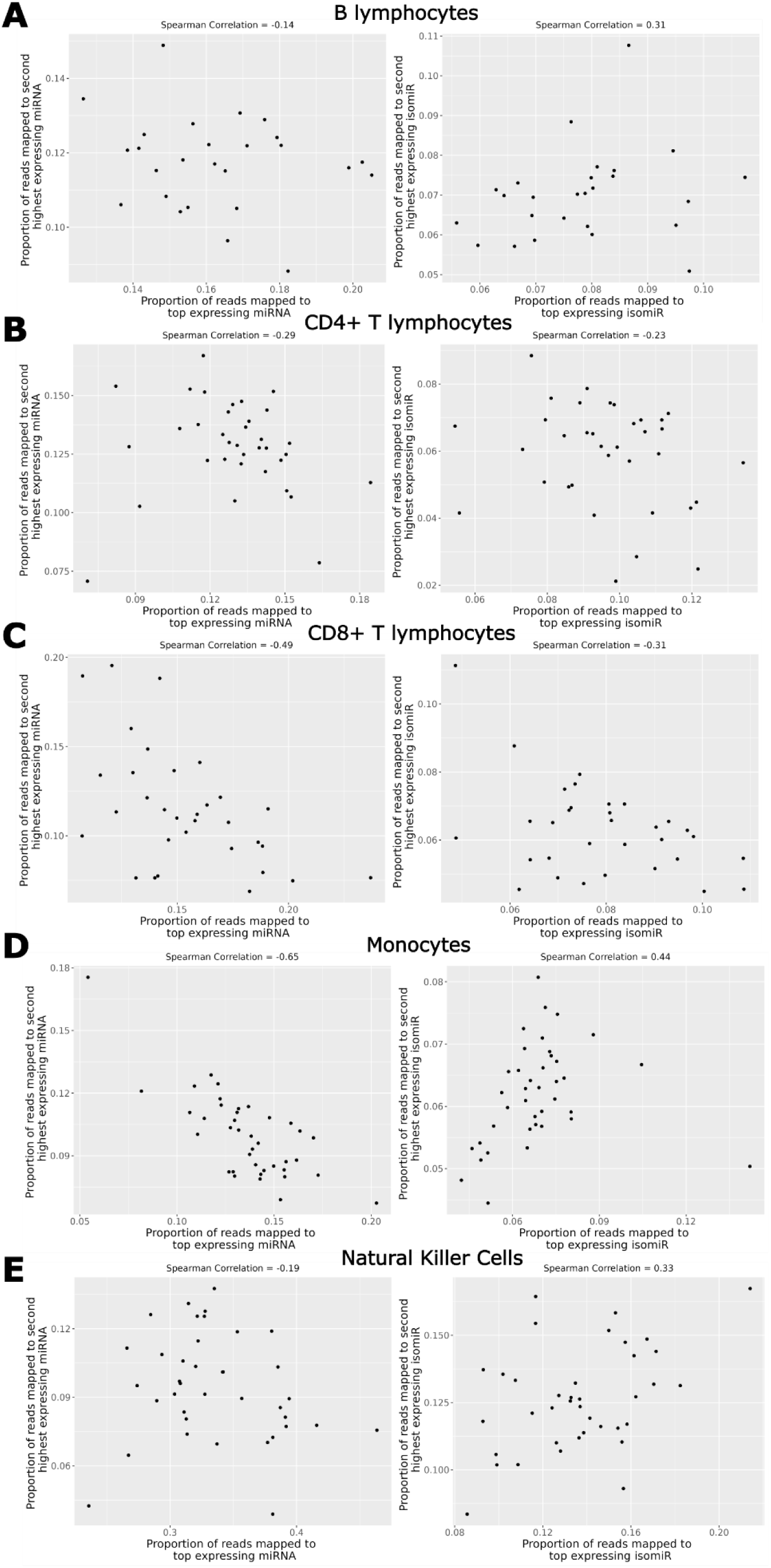
Negative correlation bias is observed when considering the top two highest expressing miRNA after aggregation to miRNA-level counts (left column, all panels), whereas a mix of positive (right column, panels A, D and E) and negative correlations (right column, panels B and C) are observed when considering the two highest expressing isomiRs.

**Supplemental Table 3:** MSE summary statistics across 100 simulations.

| method | mean | SD | median | minimum | maximum | number of<br>times<br>minimizing<br>MSE |
| --- | --- | --- | --- | --- | --- | --- |
| miRglmm | 0.01485 | 0.00463 | 0.01360 | 0.00784 | 0.03292 | 97 |
| NB GLM | 0.02315 | 0.00612 | 0.02239 | 0.01326 | 0.04883 | 0 |
| DESeq2 | 0.02192 | 0.00511 | 0.02153 | 0.01229 | 0.03608 | 3 |
| edgeR | 0.02315 | 0.00612 | 0.02239 | 0.01325 | 0.04883 | 0 |
| limmavoom | 0.02310 | 0.00622 | 0.02234 | 0.01381 | 0.04537 | 0 |

**Supplemental Table 4:** Confidence interval coverage proportion summary statistics across 100 simulations.

| method | mean | SD | median | minimum | maximum |
| --- | --- | --- | --- | --- | --- |
| miRglmm | 0.9143 | 0.0559 | 0.9262 | 0.7131 | 0.9836 |
| NB GLM | 0.7727 | 0.0641 | 0.7869 | 0.5328 | 0.8852 |
| DESeq2 | 0.8048 | 0.0449 | 0.8074 | 0.6557 | 0.8852 |
| limmavoom | 0.7869 | 0.0669 | 0.7992 | 0.5492 | 0.8934 |

**Supplemental Figure 2:**
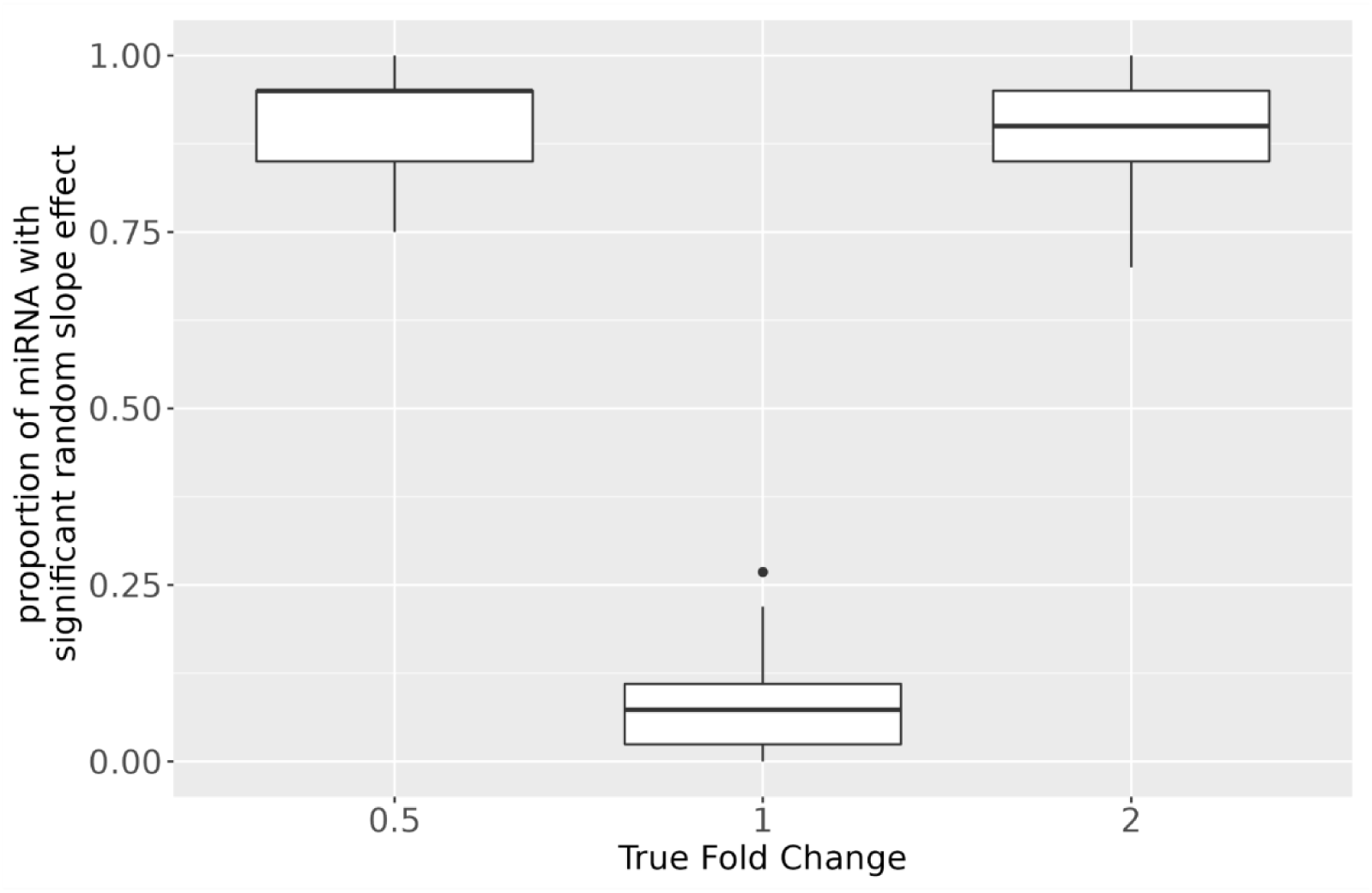
The simulation procedure creates significant variability in the group effect across isomiRs.

**Supplemental Table 5:** Comparing performance by miRglmm filters.

| method | MSE ($\times 10^{-3}$) | number of<br>miRNA<br>estimated | minimum<br>number<br>of isomiRs | maximum<br>number<br>of isomiRs | coverage<br>proportion |
| --- | --- | --- | --- | --- | --- |
| miRglmm filter -1* | 7.81 | 303 | 7 | 40 | 0.98 |
| miRglmm filter -0.5 | 7.80 | 303 | 6 | 33 | 0.97 |
| miRglmm filter 0 | 8.45 | 303 | 4 | 30 | 0.97 |
| miRglmm filter 0.5 | 8.71 | 302 | 2 | 24 | 0.98 |
| miRglmm filter 1 | 8.58 | 301 | 2 | 22 | 0.98 |
| miRglmm filter 1.5 | 9.05 | 301 | 2 | 18 | 0.99 |
| miRglmm filter 2 | 10.31 | 300 | 2 | 15 | 0.99 |
| miRglmm no filter | 15.34 | 303 | 12 | 263 | 0.97 |

MSE: Mean Squared Error, NA: not applicable, \*denotes default filter implemented in miRglmm

**Supplemental Figure 3:**
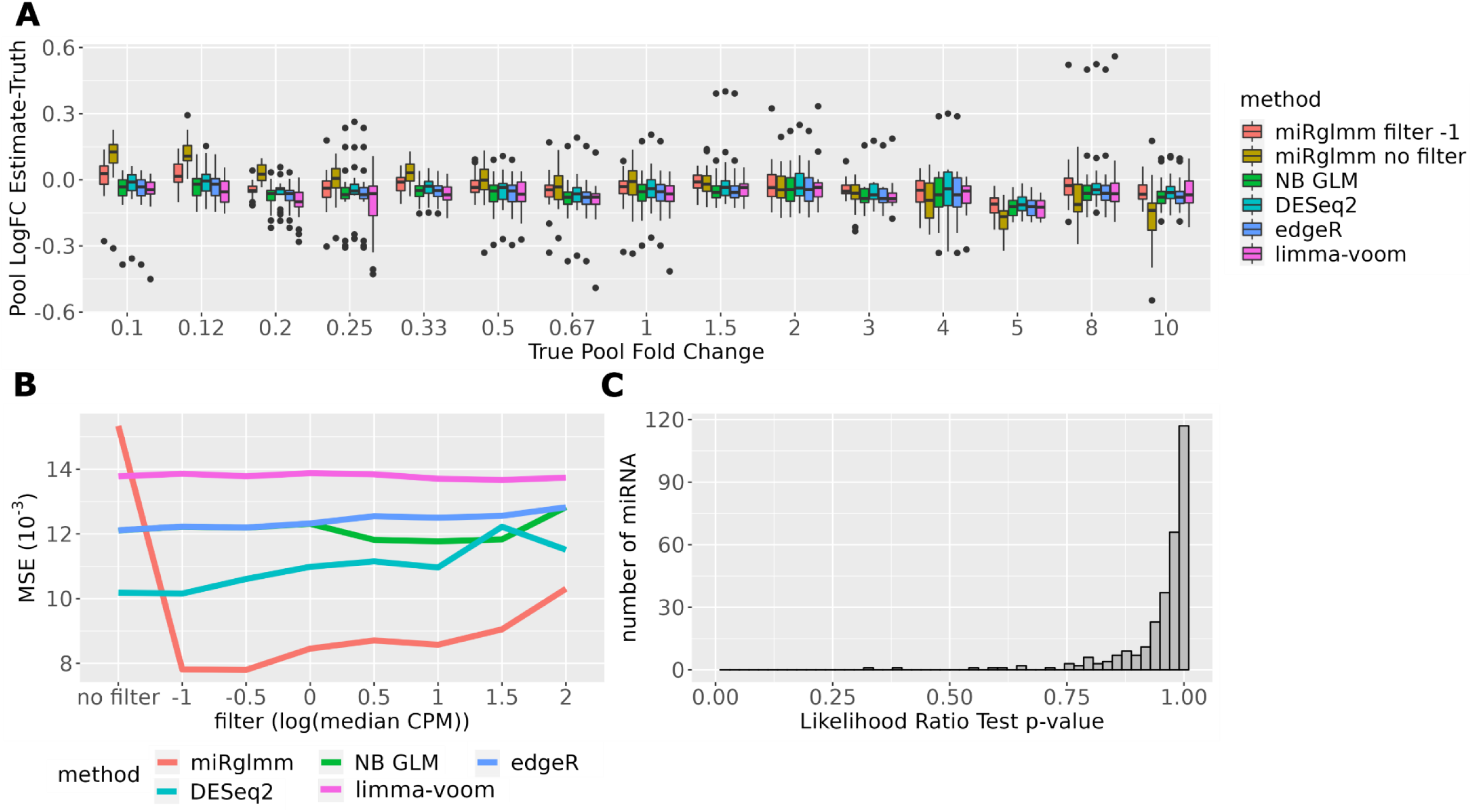
Log Fold Change (logFC) estimates of Pool B expression compared to Pool A expression are compared to known true values for each miRNA and are summarized within each True Pool Fold Change group by method (Panel B). miRglmm after filtering out isomiRs with log(median CPM < -1), miRglmm with no filtering and 4 aggregation methods (with no filtering) were used. Filtering results in a dramatic decrease in Mean Squared Error (MSE) when using miRglmm, but there is little change in performance after filtering for the aggregation methods (panel B). There are no miRNA with significant variability in the Pool effect across isomiRs as measured via a Likelihood Ratio Test p-value (panel C).

**Supplemental Figure 4:**
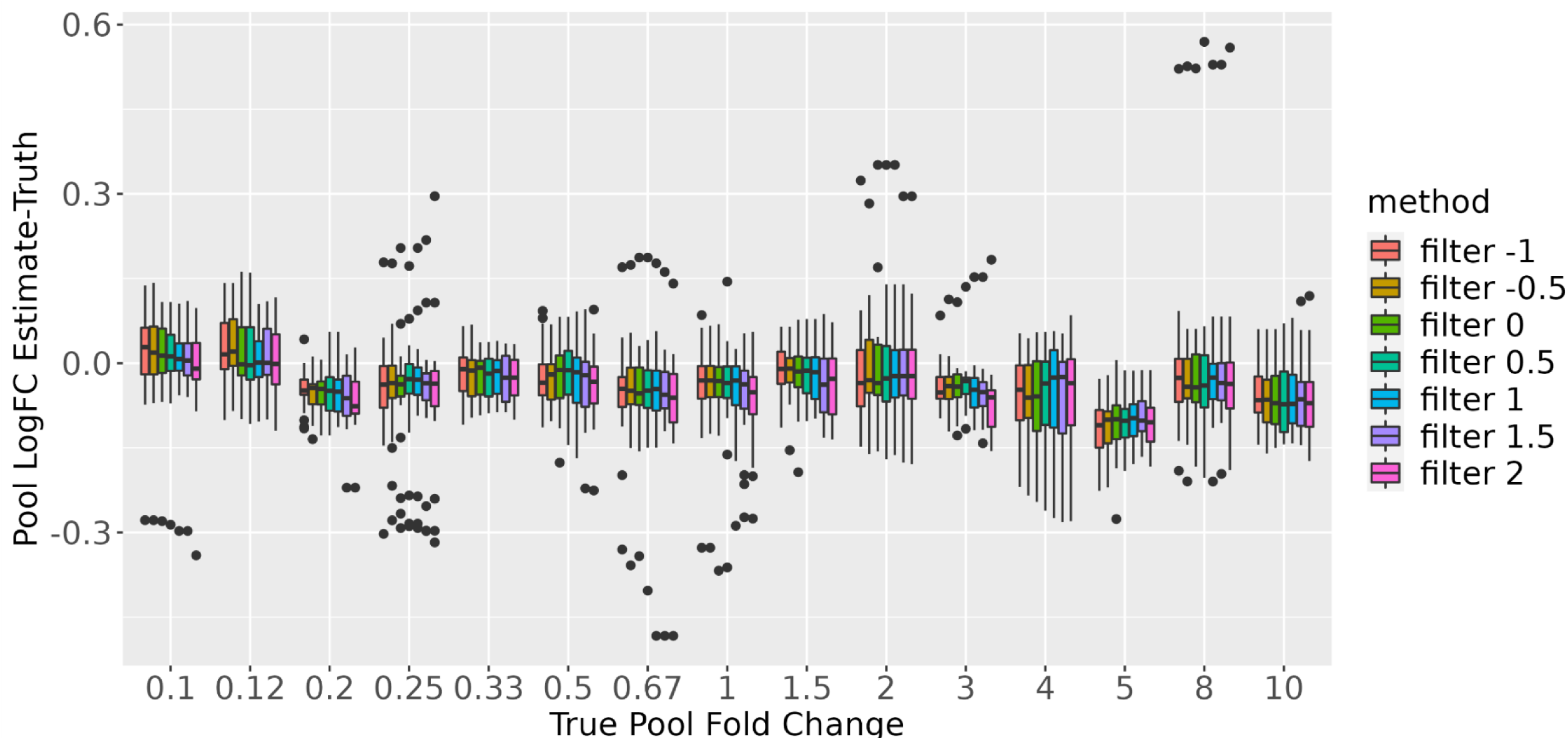
The performance of miRglmm after a variety of isomiR filters based on log(median CPM) expression appears fairly uniform. Log fold change (logFC) estimates of Pool B expression compared to Pool A expression are compared to known true values for each miRNA and summarized within True Pool Fold Change group.

**Supplemental Table 6:**
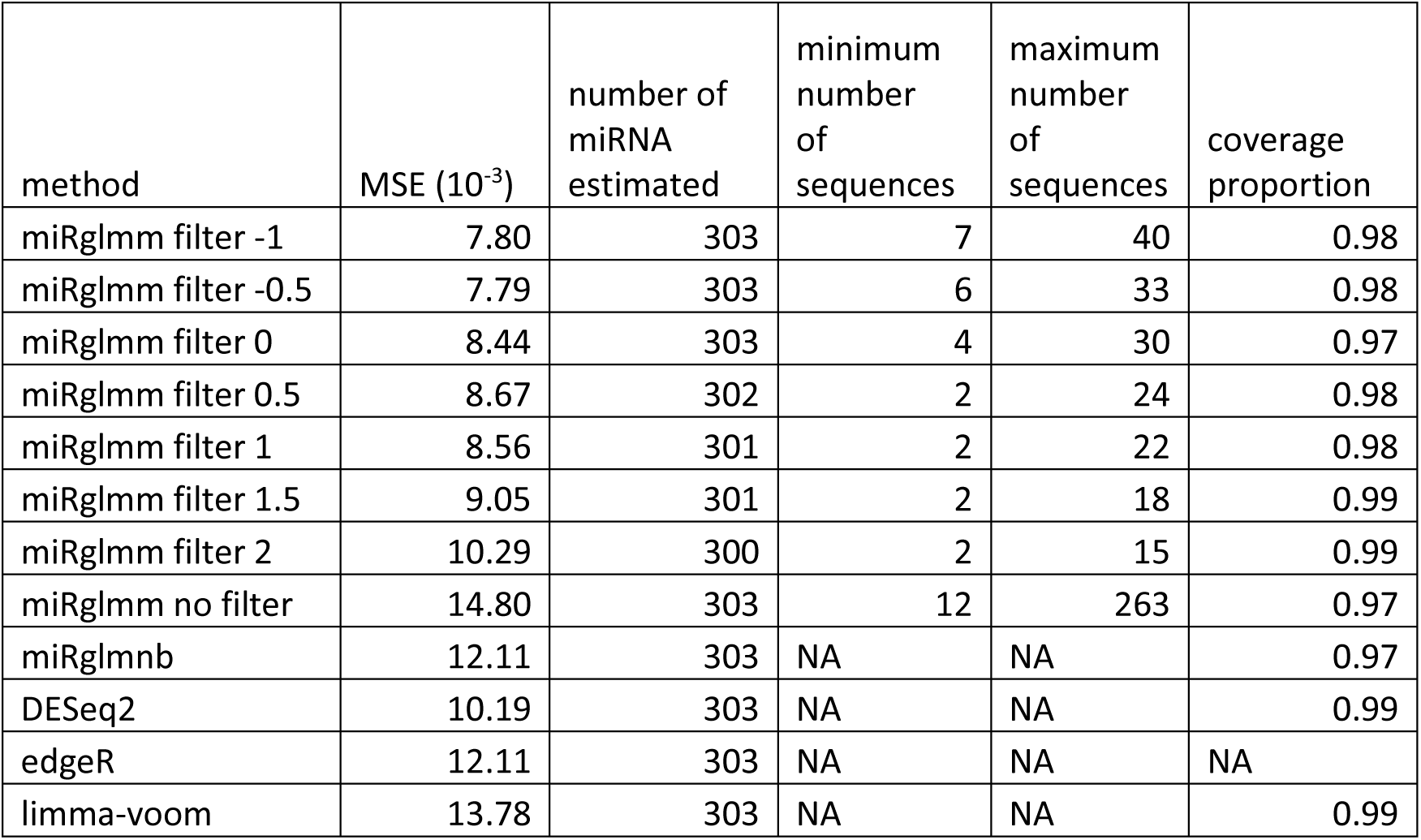
Comparing performance by miRglmm filters and comparing to aggregation methods when random slope effect removed.

**Supplemental Figure 5:**
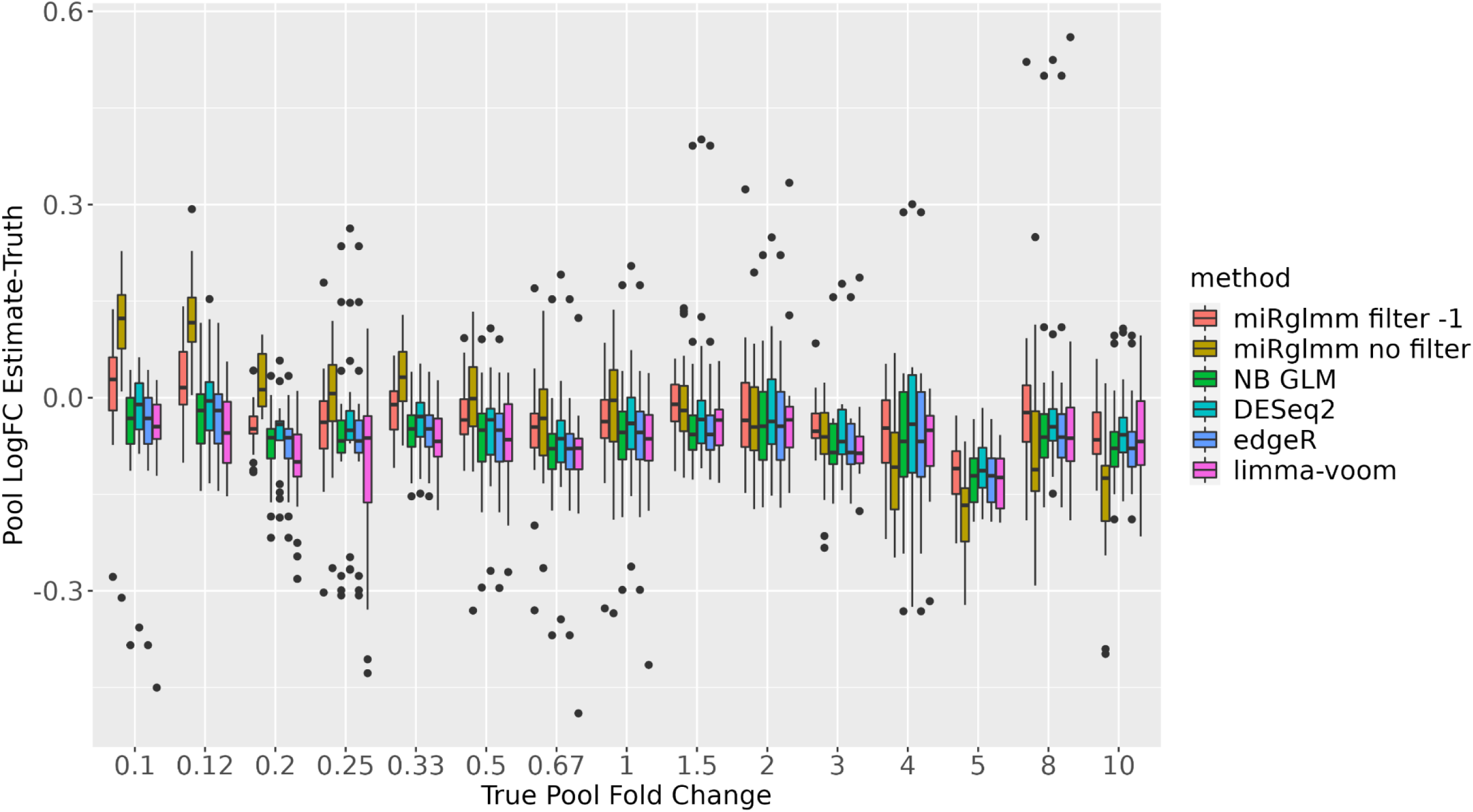
The performance of miRglmm without the random Pool effect for isomiR is compared to the performance of aggregation methods. Log fold change (logFC) estimates of Pool B expression compared to Pool A expression are compared to known true values for each miRNA and summarized within each True Pool Fold Change group.

**Supplemental Figure 6:**
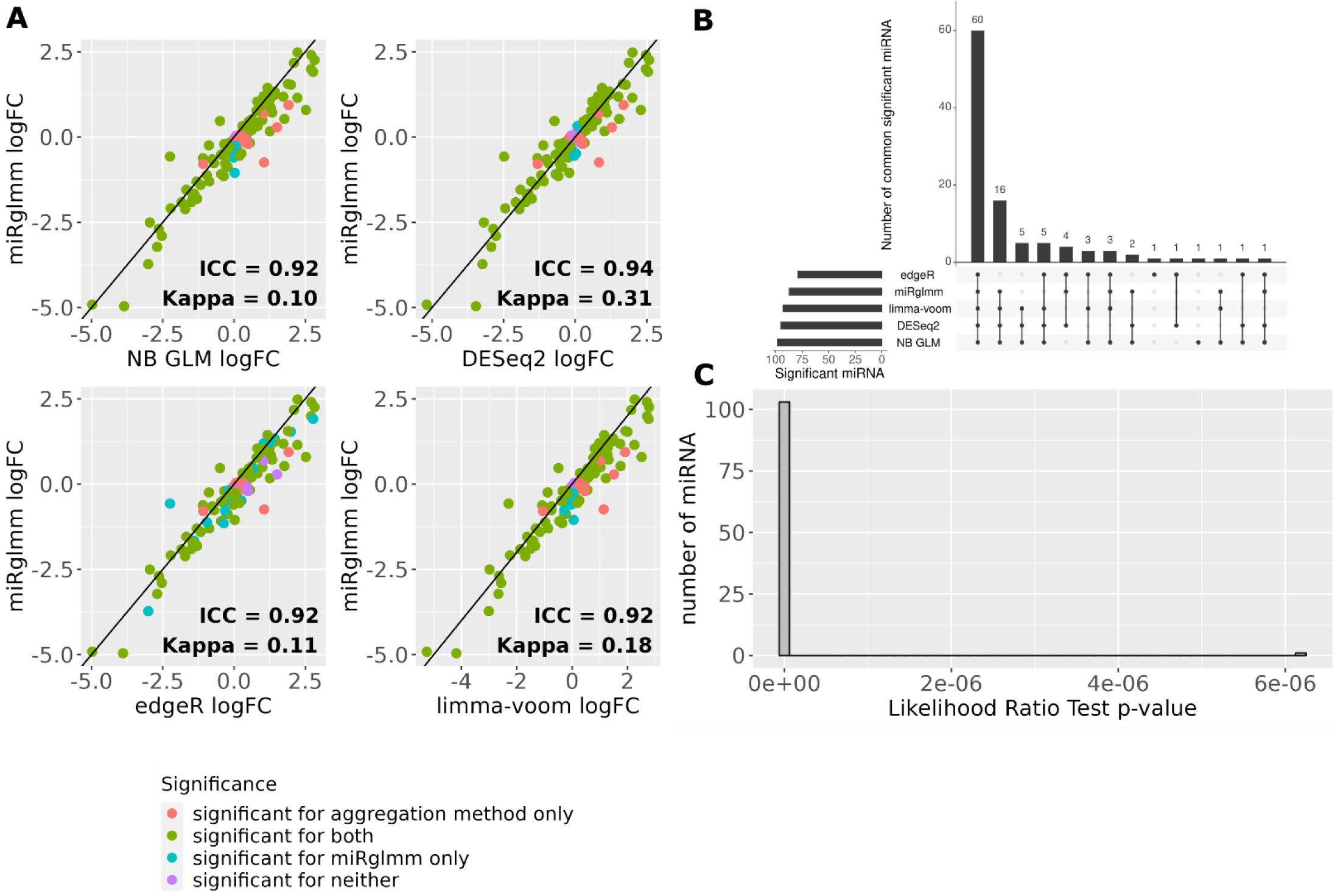
Log fold change (logFC) estimates of miRNA-level expression in monocytes compared to B lymphocytes from miRglmm are plotted versus logFC estimates from each aggregation method (panel A). Each point represents the estimate for the monocyte vs B lymphocyte contrast for a unique miRNA. Significant DE vs B lymphocytes is indicated by the color (Benjamini Hochberg FDR of 0.05). Intra-class correlation coefficients (ICC) are used to assess agreement in the values of the estimated logFC between methods. Cohen’s kappa is used to assess agreement in the significance of the estimate between methods. An upset plot shows the common number of miRNAs differentially expressed across the 5 methods (panel B). A histogram of the p-values of an LRT testing the significance of the random isomiR cell-type effect indicates that most of the miRNA have significant variability in the cell-type effect across isomiRs (panel C).

**Supplemental Figure 7:**
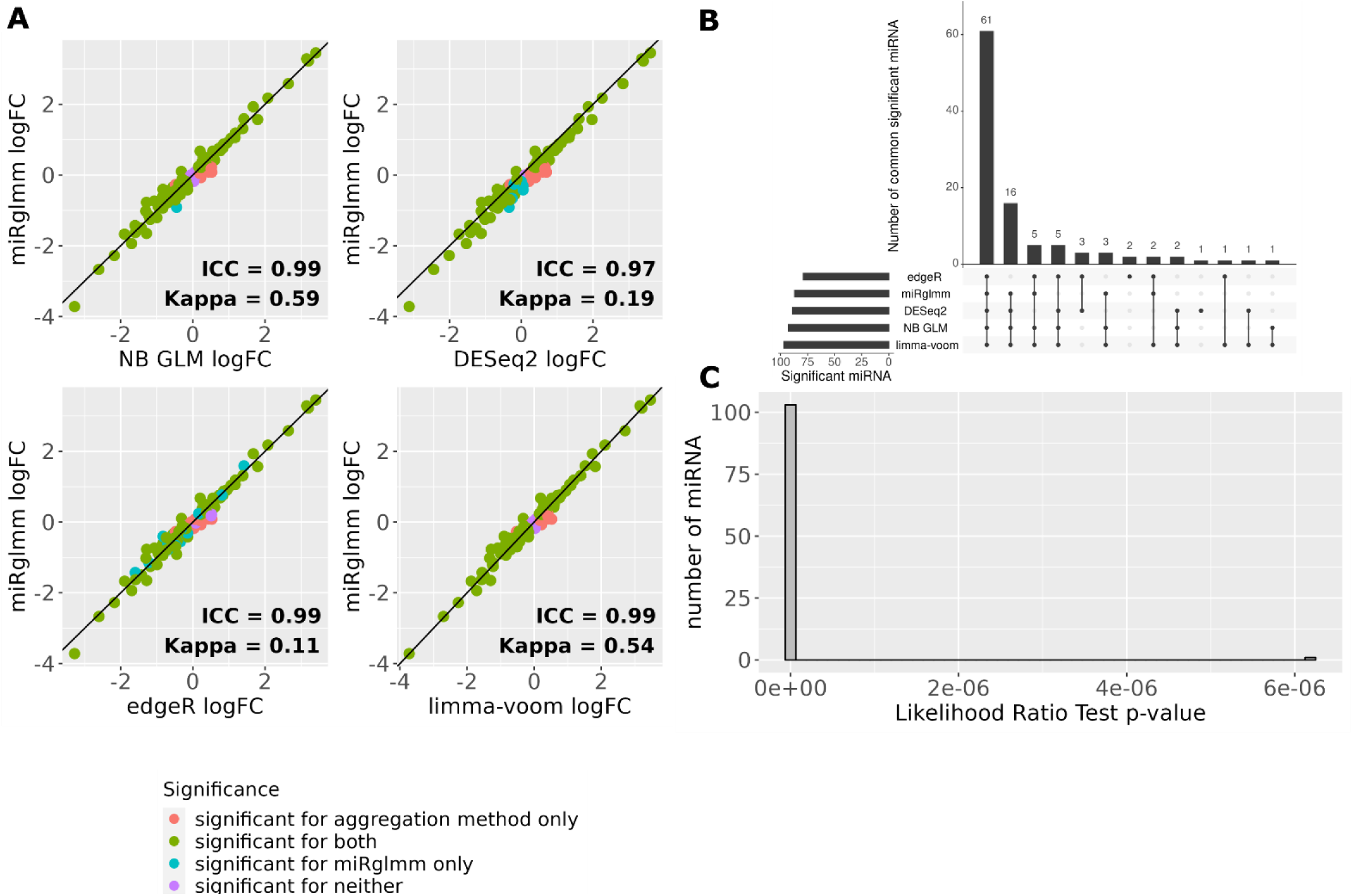
Log fold change (logFC) estimates of miRNA-level expression in Natural Killer (NK) cells compared to B lymphocytes from miRglmm are plotted versus logFC estimates from each aggregation method (panel A). Each point represents the estimate for the NK vs B lymphocyte contrast for a unique miRNA. Significant DE vs B lymphocytes is indicated by the color (Benjamini Hochberg FDR of 0.05). Intra-class correlation coefficients (ICC) are used to assess agreement in the values of the estimated logFC between methods. Cohen’s kappa is used to assess agreement in the significance of the estimate between methods. An upset plot shows the common number of miRNAs differentially expressed across the 5 methods (panel B). A histogram of the p-values of an LRT testing the significance of the random isomiR cell-type effect indicates that most of the miRNA have significant variability in the cell-type effect across isomiRs (panel C).

**Supplemental Figure 8:**
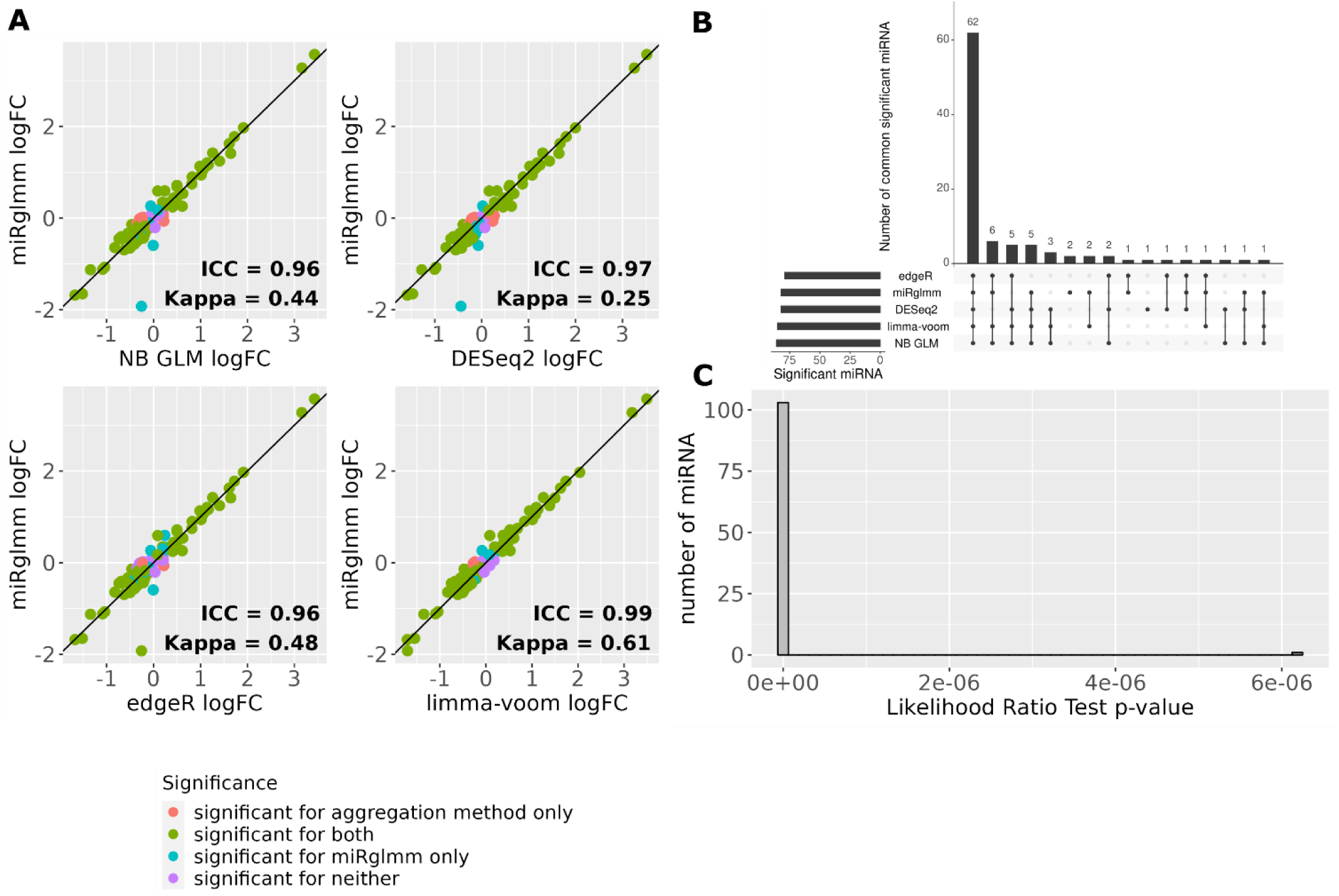
Log fold change (logFC) estimates of miRNA-level expression in CD4+ T lymphocytes compared to B lymphocytes from miRglmm are plotted versus logFC estimates from each aggregation method (panel A). Each point represents the estimate for the CD4+ T vs B lymphocyte contrast for a unique miRNA. Significant DE vs B lymphocytes is indicated by the color (Benjamini Hochberg FDR of 0.05). Intra-class correlation coefficients (ICC) are used to assess agreement in the values of the estimated logFC between methods. Cohen’s kappa is used to assess agreement in the significance of the estimate between methods. An upset plot shows the common number of miRNAs differentially expressed across the 5 methods (panel B). A histogram of the p-values of an LRT testing the significance of the random isomiR cell-type effect indicates that most of the miRNA have significant variability in the cell-type effect across isomiRs (panel C).

**Supplemental Figure 9:**
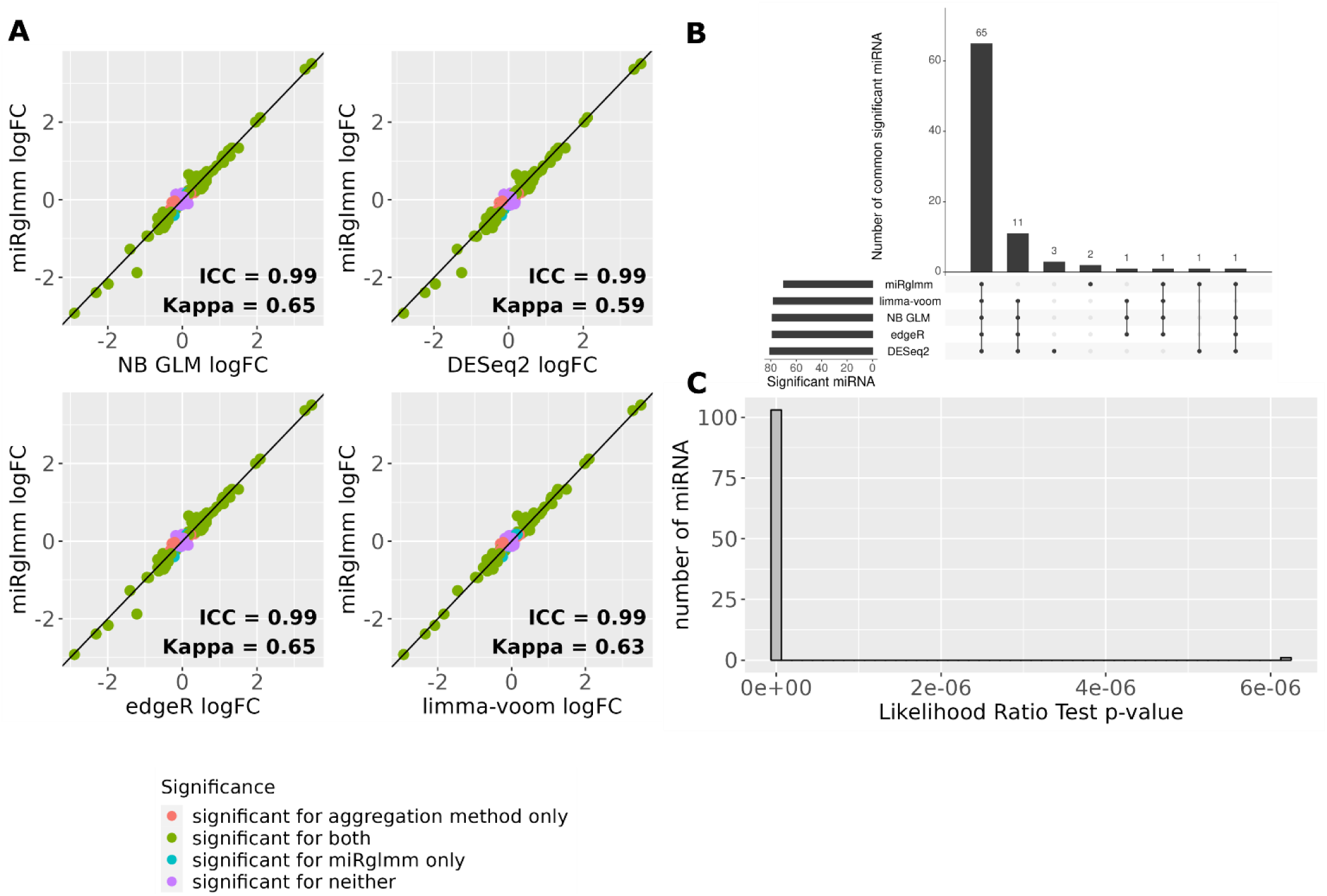
Log fold change (logFC) estimates of miRNA-level expression in CD8+ T lymphocytes compared to B lymphocytes from miRglmm are plotted versus logFC estimates from each aggregation method (panel A). Each point represents the estimate for the CD8+ T vs B lymphocyte contrast for a unique miRNA. Significant DE vs B lymphocytes is indicated by the color (Benjamini Hochberg FDR of 0.05). Intra-class correlation coefficients (ICC) are used to assess agreement in the values of the estimated logFC between methods. Cohen’s kappa is used to assess agreement in the significance of the estimate between methods. An upset plot shows the common number of miRNAs differentially expressed across the 5 methods (panel B). A histogram of the p-values of an LRT testing the significance of the random isomiR cell-type effect indicates that most of the miRNA have significant variability in the cell-type effect across isomiRs (panel C).

**Supplemental Table 7:** LogFC estimates for miRNA-contrast pairs that are significant for miRglmm but not significant for any aggregation method.

| miRNA | Contrast (vs B lymphocyte) | miRglmm | NB GLM | DESeq2 | edgeR | limma-voom |
| --- | --- | --- | --- | --- | --- | --- |
| hsa-miR-32-5p | CD4 T lymphocyte | 0.26 | -0.06 | 0.02 | -0.06 | -0.08 |
| hsa-miR-340-5p | CD8 T lymphocyte | -0.4 | -0.22 | -0.19 | -0.22 | -0.25 |
| hsa-miR-423-3p | CD8 T lymphocyte | -0.15 | -0.06 | 0 | -0.06 | -0.07 |
| hsa-miR-423-5p | CD4 T lymphocyte | -0.38 | -0.25 | -0.15 | -0.25 | -0.26 |

**Supplemental Table 8:** LogFC estimates for miRNA-contrast pairs that are not significant for miRglmm but significant for all aggregation methods.

| miRNA | contrast | miRglmm | NB GLM | DESeq2 | edgeR | limma-voom |
| --- | --- | --- | --- | --- | --- | --- |
| hsa-let-7b-5p | Monocyte | 0.09 | 0.45 | 0.22 | 0.45 | 0.47 |
| hsa-miR-103a-3p | CD8 T lymphocyte | 0.14 | 0.17 | 0.21 | 0.17 | 0.16 |
| hsa-miR-106b-3p | CD8 T lymphocyte | -0.07 | -0.28 | -0.25 | -0.28 | -0.3 |
| hsa-miR-107 | Natural Killer | -0.07 | 0.22 | 0.39 | 0.22 | 0.22 |
| hsa-miR-107 | CD8 T lymphocyte | 0.12 | 0.17 | 0.21 | 0.17 | 0.16 |
| hsa-miR-140-3p | CD4 T lymphocyte | -0.1 | -0.19 | -0.12 | -0.19 | -0.18 |
| hsa-miR-140-3p | CD8 T lymphocyte | -0.12 | -0.22 | -0.17 | -0.22 | -0.21 |
| hsa-miR-148b-3p | Monocyte | 0.94 | 1.91 | 1.69 | 1.91 | 1.92 |
| hsa-miR-148b-3p | CD8 T lymphocyte | 0.18 | 0.32 | 0.34 | 0.32 | 0.26 |
| hsa-miR-181a-3p | Monocyte | -0.75 | 1.05 | 0.83 | 1.05 | 1.14 |
| hsa-miR-181a-3p | CD4 T lymphocyte | -0.13 | -0.28 | -0.22 | -0.28 | -0.23 |
| hsa-miR-181a-3p | CD8 T lymphocyte | -0.06 | -0.27 | -0.2 | -0.27 | -0.19 |
| hsa-miR-181b-5p | CD4 T lymphocyte | 0 | -0.28 | -0.2 | -0.28 | -0.23 |
| hsa-miR-221-3p | Natural Killer | -0.27 | -0.48 | -0.31 | -0.48 | -0.53 |
| hsa-miR-26b-3p | CD8 T lymphocyte | 0.17 | 0.22 | 0.26 | 0.22 | 0.21 |
| hsa-miR-28-5p | CD8 T lymphocyte | -0.05 | -0.15 | -0.11 | -0.15 | -0.17 |
| hsa-miR-30d-5p | Monocyte | -0.04 | 0.33 | 0.1 | 0.33 | 0.33 |
| hsa-miR-30d-5p | CD4 T lymphocyte | -0.07 | -0.2 | -0.13 | -0.2 | -0.19 |
| hsa-miR-320a-3p | CD8 T lymphocyte | 0.11 | 0.14 | 0.2 | 0.14 | 0.13 |
| hsa-miR-339-3p | Monocyte | -0.79 | -1.08 | -1.31 | -1.08 | -1.06 |
| hsa-miR-374a-3p | Natural Killer | 0.09 | 0.43 | 0.6 | 0.43 | 0.4 |
| hsa-miR-374a-5p | Natural Killer | 0.05 | 0.19 | 0.36 | 0.19 | 0.18 |
| hsa-miR-378a-3p | CD8 T lymphocyte | -0.23 | -0.32 | -0.27 | -0.32 | -0.3 |
| hsa-miR-652-3p | CD4 T lymphocyte | -0.01 | -0.23 | -0.19 | -0.23 | -0.28 |
| hsa-miR-652-3p | CD8 T lymphocyte | -0.03 | -0.21 | -0.17 | -0.21 | -0.23 |
| hsa-miR-941 | Natural Killer | 0.08 | 0.52 | 0.7 | 0.52 | 0.52 |

**Supplemental Figure 10:**
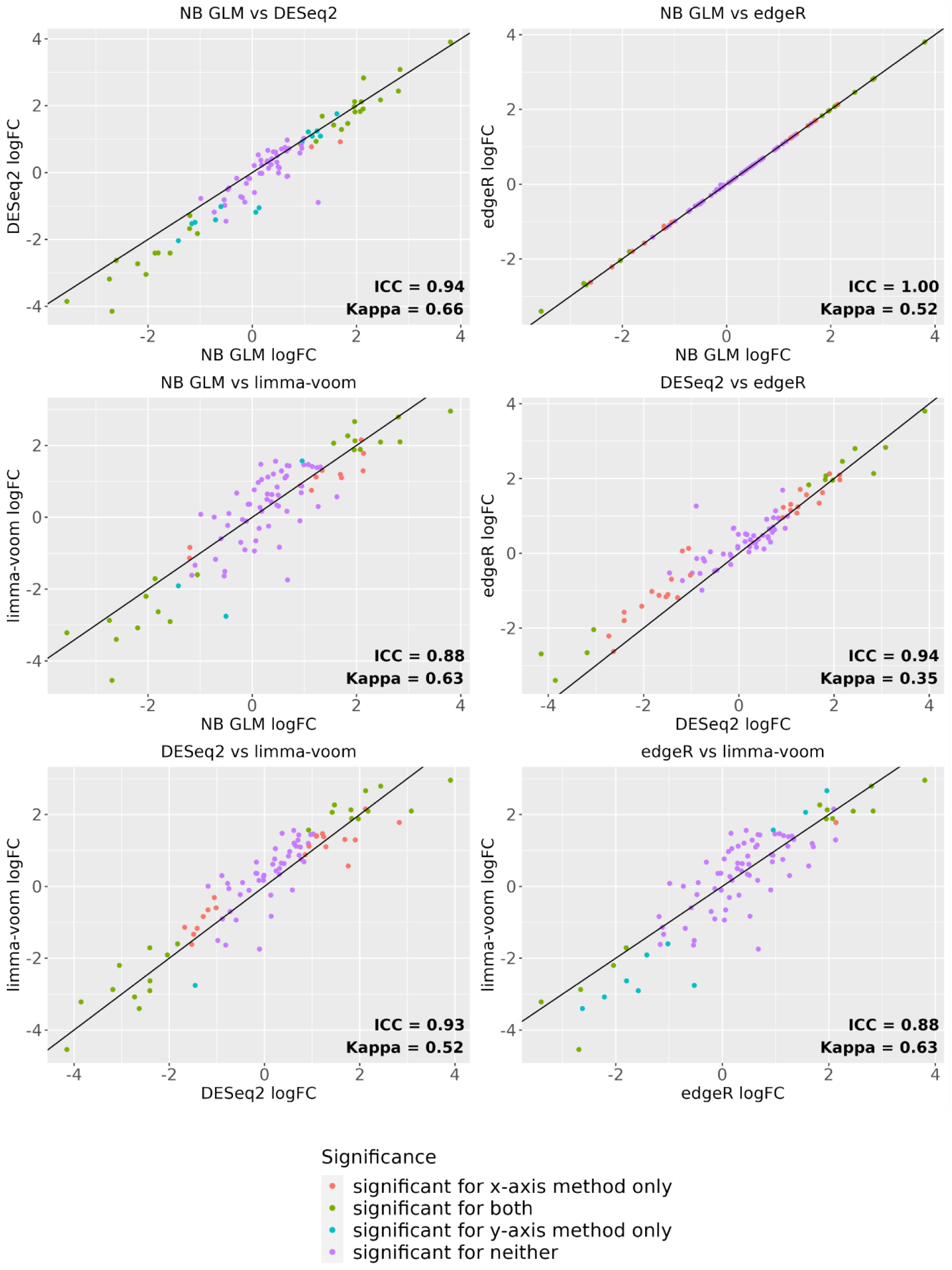
Log fold change (logFC) estimates of expression in testes compared to bladder are compared between aggregation methods with differences in significance (Benjamini Hochberg FDR of 0.05) indicated by color.

**Supplemental Figure 11:**
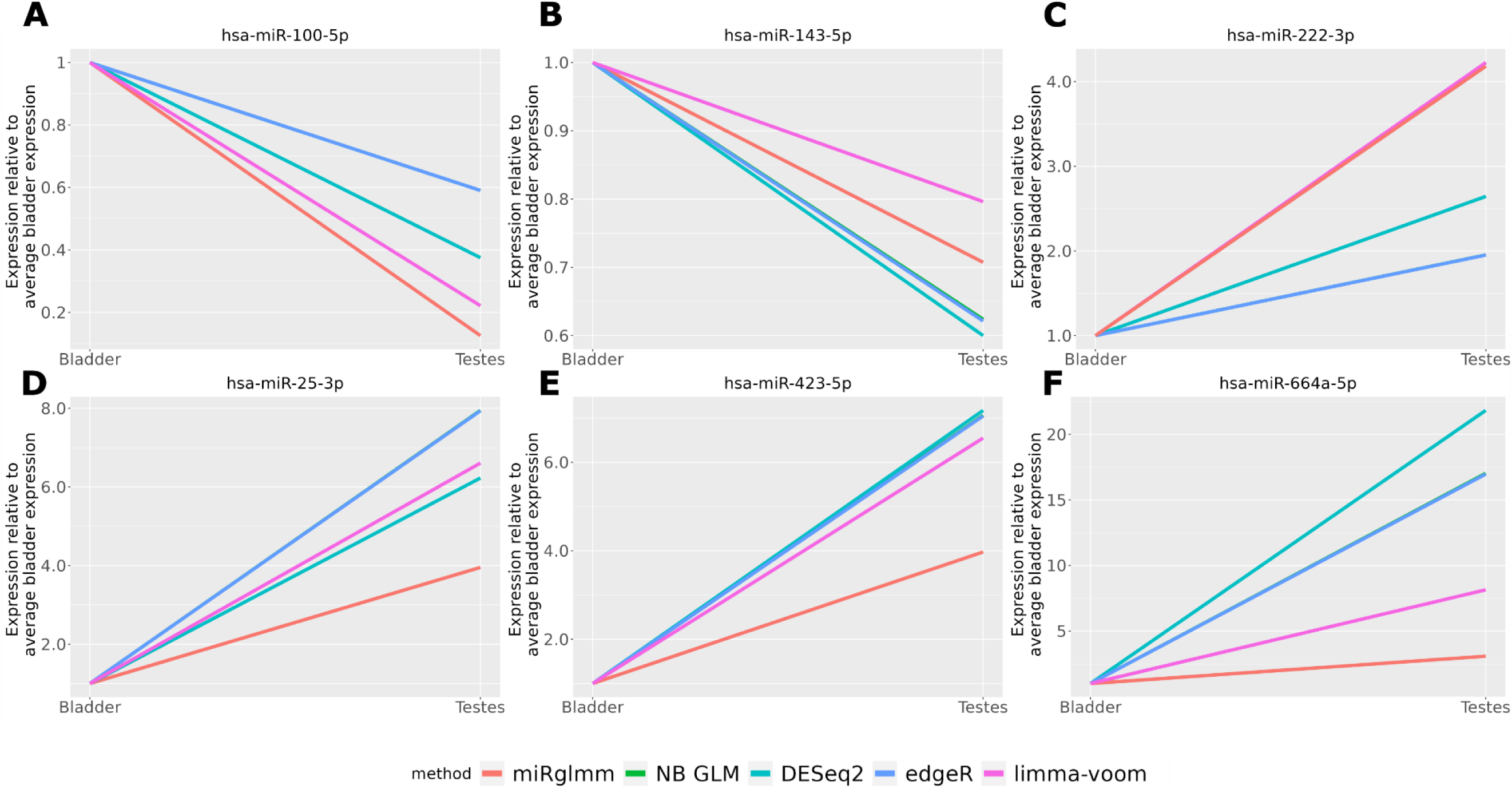
Estimates of testes expression relative to bladder are compared between miRglmm and all aggregation methods. miRglmm and limma-voom provide nearly identical estimates for hsa-miR-222-3p. NB GLM and edgeR provide nearly identical estimates for all miRNA.

**Supplemental Figure 12:**
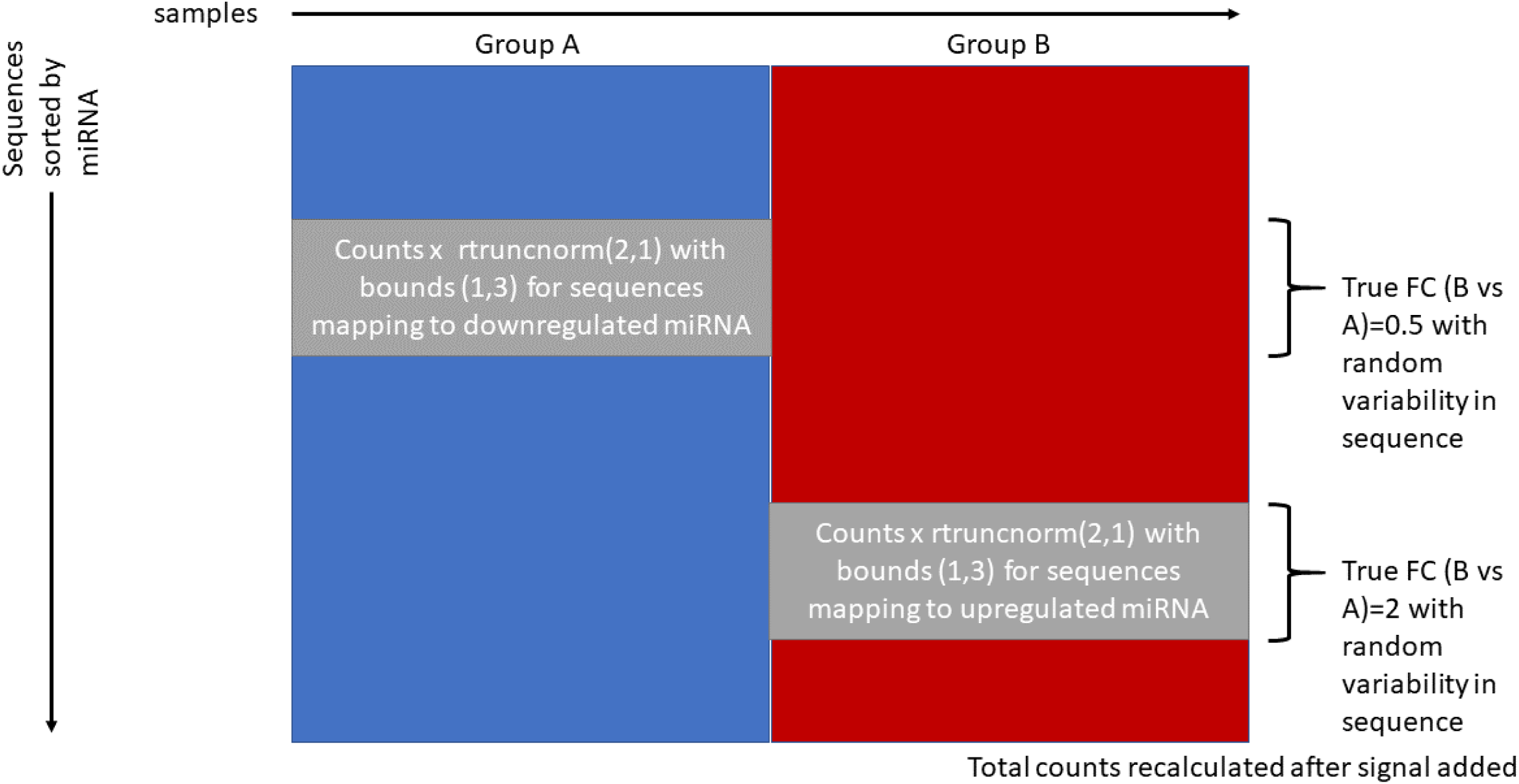
Inducing an artificial group effect with random sequence variability into monocyte miRNA-seq data. The entire procedure from random splitting of samples into groups is repeated 100 times.

**Supplemental Figure 13:**
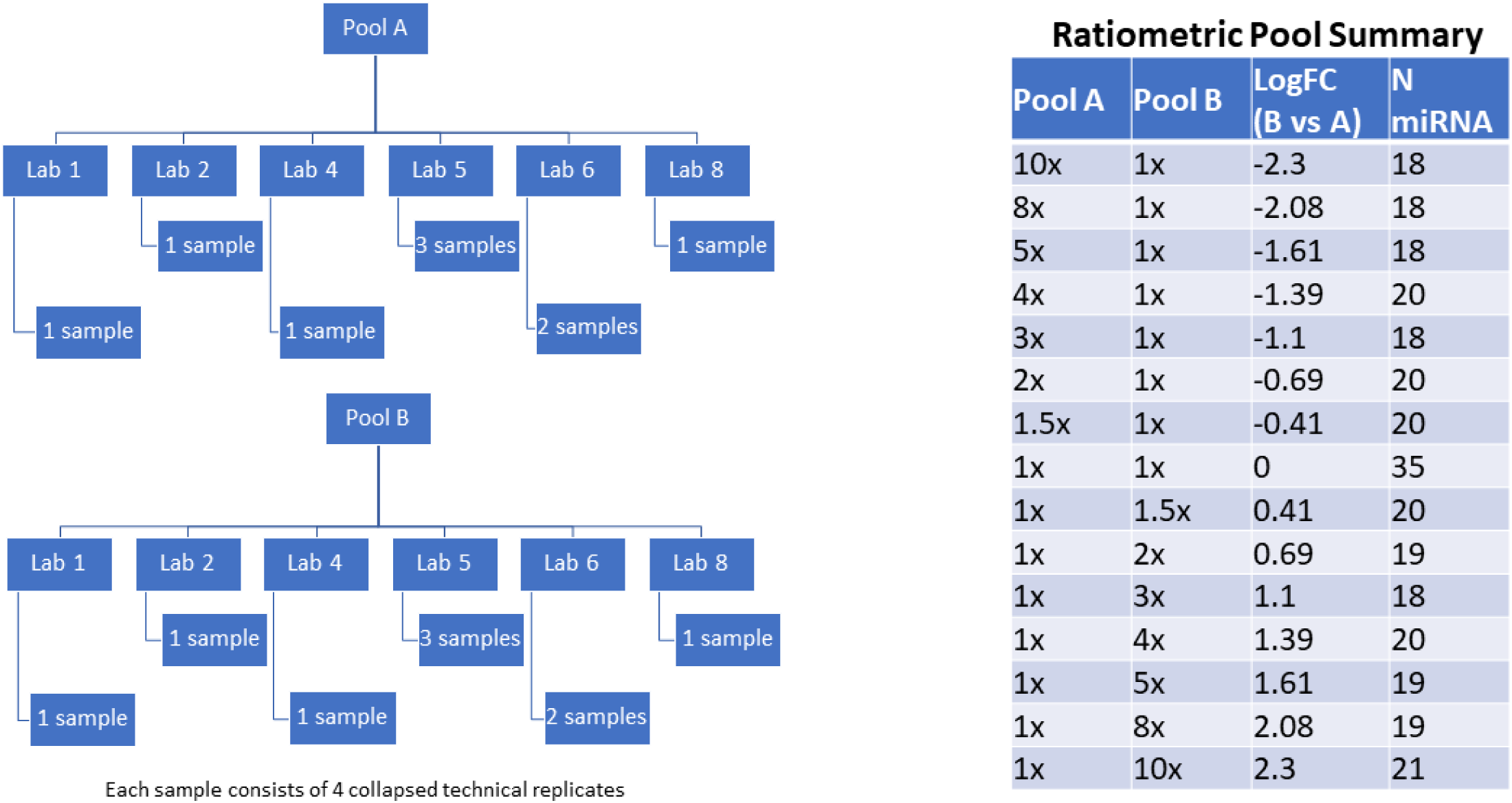
ERCC experimental design. Samples from two pools were generated by six laboratories. Each synthetic miRNA was added to Pool A and Pool B at one of fifteen possible ratios.

**Supplemental Figure 14:**
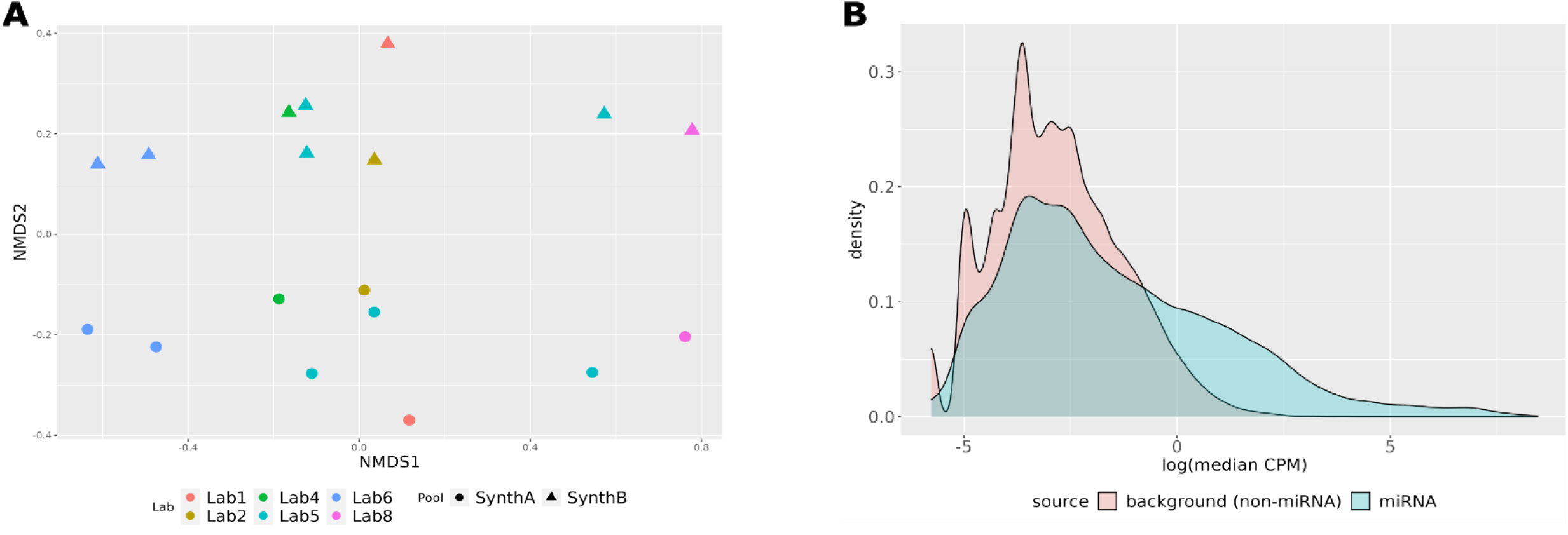
Non-metric multidimensional rescaling (NMDS) of raw counts reveals separation of samples based on laboratory along the first dimension and separation based on Pool along the second dimension (panel A). Density plots comparing the distributions of expression between sequences mapped to miRNA and sequences not mapped to miRNA can be used to define a threshold for retaining sequences with expression above background levels (panel B).

